# Wide-ranging behavioral dysfunction in two mouse models of pathological human variants in the *GRIK2* kainate receptor gene

**DOI:** 10.1101/2025.08.08.667766

**Authors:** Brynna T. Webb, Hieu Trinh, Emily A. Breach, Kendall M. Foote, Erica Binelli, Geoffrey T. Swanson

## Abstract

*De novo* variants in a subset of ionotropic glutamate receptor (iGluR) genes cause nonsyndromic neurodevelopmental disorders (NDDs) in individuals. Two recurrent variants in the kainate receptor (KAR) gene *GRIK2* result in the gain-of-function (GoF) substitutions p.Ala657Thr and p.Thr660Lys in a critical pore-forming domain of the GluK2 subunit. Disorders in individuals with these variants manifest as intellectual disability, developmental delay, motor impairments, and, in the case of p.Thr660Lys, epilepsy. To explore their pathogenicity and phenotypic consequences *in vivo*, we generated knock-in mouse models harboring orthologous *Grik2* mutations. Behavioral analyses revealed a range of developmental, motor, cognitive, and naturalistic behavior impairments in both lines, with T660K mice typically exhibiting more severe phenotypes, consistent with clinical observations in humans. GluK2(T660K) mice also display interictal EEG abnormalities and handling-induced seizures. These models establish the first *in vivo* platforms for dissecting the underlying mechanisms of NDDs caused by a GoF mutation in the GluK2 KAR subunit and represent crucial tools for therapeutic development.

**Highlights:**

- *De novo* variants in the *GRIK2* kainate receptor gene cause developmental disorders
- We studied knockin mouse models of pathogenic p.Ala657Thr and p.Thr660Lys variants
- GluK2(A657T) and GluK2(T660K) mice have deficits analogous to human disorder symptoms
- Like their human counterparts, GluK2(T660K) but not GluK2(A657T) mice have seizures
- *Grik2* knockin mice are key tools for interrogating underlying causes of dysfunction

## 1. Introduction

Developmental disorders affect approximately 15% of the pediatric population, making them among the most common conditions impacting the lives of children (Zablotsky et al., 2019). Neurodevelopmental disorders (NDDs) have wide ranging etiologies and, while some fit into specific syndromes with well-characterized spectrums of symptoms, many fall outside of these characterizations as non-syndromic NDDs. Monogenic etiologies are estimated to contribute to the incidence of NDDs at a rate of approximately 329 per 100,000 births, which equates to over 400,000 new cases every year (de Ligt et al., 2013; Lopez-Rivera et al., 2020), and are frequently identified as potential causative mutations through clinical genetic testing (Wilfert et al., 2017). Deleterious mutations in ionotropic glutamate receptor (iGluR) genes encoding AMPA (a-amino-3-hydroxy-5-methyl-4-isoxazolepropionic acid), NMDA (N-methyl-D-aspartic acid), kainate, or delta receptors are causative for numerous non-syndromic NDDs (Hansen et al., 2021).

Kainate receptors (KARs) are glutamate-gated ion channels expressed throughout the mammalian central nervous system (CNS) that modulate cellular excitability, shape pre- and postsynaptic neurotransmission, and influence synaptic development (Contractor et al., 2011; Hansen et al., 2021; Mulle and Crepel, 2021). The KAR subunit proteins are encoded by the *GRIK1-5* genes, and functional receptors are formed from tetrameric assemblies of the obligate GluK1, −2, or −3 subunits together with GluK4 or GluK5 subunits (Hansen et al., 2021). Three KAR genes, *GRIK2, GRIK3,* and *GRIK5*, are intolerant to loss-of-function (LoF) and nonsynonymous single nucleotide variants (SNVs) in the human population (Karczewski et al., 2020).

Disruption of the *GRIK2* KAR gene can be causative for ultrarare NDDs. Missense variants in *GRIK2* underlie NDDs of varying degrees of severity in patients with intellectual disability (ID), global developmental delay (GDD), impairments in the use of language, and, in a subset of patients, epilepsy (Guzmán et al., 2017; Stolz et al., 2021). As well, inherited homozygous protein truncating variants (PTVs) were proposed to underlie NDDs in siblings from a consanguineous family (Córdoba et al., 2015) as were inherited complex gene deletions that included *GRIK2* (Motazacker et al., 2007). Earlier studies using linkage disequilibrium and genome-wide association studies also implicated *GRIK2* in autism spectrum disorders (ASDs) (Jamain et al., 2002; Kim et al., 2007; Shuang et al., 2004), and neuropsychiatric disorders (Bah et al., 2004; Delorme et al., 2004; Ma et al., 2019; Mattheisen et al., 2015; Sampaio et al., 2011).

The most common damaging *GRIK2* SNV, c.1969G>A, occurs in nine individuals and changes alanine 657 to a threonine (p.Ala657Thr) in the pore-forming M3 domain of the GluK2 subunit (GenBank: NP_068775). The second most common and highly detrimental variant, c.1979C>A, has been identified in three individuals. This variant causes the substitution of threonine 660 with lysine (p.Thr660Lys) at the tip of the pore-forming M3 domain of the GluK2 subunit (GenBank: NP_068775).

Recombinant GluK2 subunits with the analogous A657T or T660K mutations form gain-of-function (GoF) KARs that exhibit a very high affinity for glutamate and a prolonged current decay in patch-clamp recordings from heterologous cell lines (Guzmán et al., 2017; Kohda et al., 2000; Stolz et al., 2021).

While both variants lead to a putative GoF in GluK2-containing KARs, their associated nonsyndromic disorders exhibit a mix of shared and distinct symptoms, reflecting their related yet distinct etiologies. All individuals with the p.Ala657Thr *de novo* variant have a suite of symptoms that are similar in nature but vary in severity, including ID, GDD, speech dysfunction, and motor dysfunction that manifest clinically as ataxia, uncoordinated gait, or imbalance. None of the individuals with the *GRIK2* p.Ala657Thr variant have seizures and their MRIs, where performed, have not revealed any gross morphological abnormalities (Guzmán et al., 2017; Stolz et al., 2021). The three individuals with the p.Thr660Lys variant suffer from profoundly debilitating symptoms – to the extent that none are able to walk or speak and exhibit no or limited responsiveness to external stimuli. All three individuals have seizure disorders varying in severity, and their MRIs reveal white matter abnormalities and reductions in neuropil volume (Guzmán et al., 2017; Stolz et al., 2021). The genetic, functional, and clinical data therefore are consistent with *GRIK2* variants causing autosomal dominant, monogenic NDDs.

We recently generated knockin mouse models of the GoF *GRIK2* p.Ala657Thr and p.Thr660Lys variants with analogous mutations in the mouse *Grik2* gene, referred to as the GluK2(A657T) and GluK2(T660K) mouse lines, respectively, to gain insight into mechanisms underlying the human NDDs. In this report, we compare heterozygous GluK2(A657T) and GluK2(T660K) mice with their wildtype (WT) littermates in a battery of behavioral tests to determine the extent to which these mutations cause dysfunction similar to that observed in humans with the *GRIK2* p.Ala657Thr and p.Thr660Lys variants. Our results suggest that these two *Grik2* mutant lines have both distinct and overlapping postural, motor, cognitive, developmental, and other behavioral deficits, supporting face validity of these models. Our analysis provides further evidence that both *GRIK2* c.1969G>A and c.1979C>A are causative for NDDs in humans. This work provides a framework for studying the role of KAR gene GoF variants in NDD and will enable future studies into the mechanisms by which KAR dysfunction causes NDDs.

## 2. Materials and Methods

### 2.1. Animals

All experimental procedures were conducted in accordance with the ethical policies and protocols approved by the Northwestern University Institutional Animal Care and Use Committee. The GluK2(A657T) and GluK2(T660K) mouse lines were created using CRISPR/Cas9 editing to introduce the orthologous *Grik2* c.1969G>A or c.1979C>A point variants in the mouse gene in the Northwestern University Transgenic and Targeted Mutagenesis (TTML) facility (Nomura et al., 2023). Mice were housed with *ad libitum* access to food and water and with a 12/12 h dark-light cycle except for food-restricted animals used for T-Maze testing. Mice of both sexes were used in all tests in approximately equal ratios. Experiments were performed by investigators blind to animal genotype. Data was uncoded following statistical analysis.

### 2.2. Genotyping

Genotyping was performed on ear or tail biopsy samples obtained during weaning. DNA was extracted from the samples and amplified via PCR using a mixture of Colorless GoTaq Master Mix (Promega Corporation, Madison, WI, USA), nuclease-free water, Grik2-Forward Primer (TTG TCA CAT GGC ACC CTC TC), and Grik2-Reverse Primer (GGT TTC AGT GGC AAG TAA GGC) (Integrated DNA Technologies, Inc., Coralville, IA, USA). For the A657T line, restriction digestion was performed on the PCR product using Buffer R (with BSA) (Thermo Fisher Scientific Inc., Waltham, MA, USA), TaiI (MaeII) restriction enzyme (Thermo Fisher Scientific Inc., Waltham, MA, USA), and nuclease-free water. DNA was electrophoresed on a 2.5% agarose gel and imaged using a LI-COR Odyssey XF (LI-COR Biotech, Lincoln, NE, USA) to determine genotypes from band sizes. For the T660K mouse line, the PCR product was purified using the Gel/PCR DNA Fragments Extraction Kit (IBI Scientific, Dubuque, IA, USA) and sequenced by the Northwestern University Sanger Sequencing Core.

### 2.3. Synaptosome preparation and Western blot

Synaptosome preparations were made as previously described (Armstrong et al., 2002; Armstrong et al., 2006). Hippocampi isolated from adult mice were homogenized in 0.325 M sucrose and then centrifuged at 900 g for 10 min at 4°C. The pellet was reserved on ice, and the supernatant was collected and centrifuged at 17,000 g for 55 min at 4°C. The resultant pellet was enriched with small hippocampal synaptosomes (P2). The large, dense pellet contained mossy fiber synapses sedimented with the nuclei and other dense cellular debris after the initial low-speed spin. This pellet was resuspended in 1.5 ml of 18% Ficoll in 0.325 M sucrose and centrifuged at 7,500 g for 40 min. The supernatant from this spin was collected, diluted in 4 volumes 0.325 M sucrose, and centrifuged at 13,000 g for 20 min at 4°C to yield a pellet containing the large mossy fiber synaptosomes (P3).

Protein samples from P2 and P3 fractions were diluted in 0.325 M sucrose, 20 mM Tris-buffered saline (TBS) and NuPAGE lithium dodecyl sulfate (LDS) loading buffer (pH 8.4), heated to 70°C for 10 min while mixing on a thermomixer. Diluted protein samples (approximately 10 micrograms) were run on a precast 4%-12% gradient polyacrylamide Bis-Tris NuPAGE gel in MOPS/SDS running buffer (50 mM MOPS, 50 mM Tris Base, 0.1% SDS, 1 mM EDTA, pH 7.7, 0.01-0.09% N,N-dimethylformamide). Separated protein was then transferred to PVDF membrane (overnight at low voltage), washed in TBS (2 x 5 minutes) blocked in TBS containing 3% IgG-free BSA and 1% normal donkey serum (>1 hour), and blotted overnight in blocking solution with the following antibodies: mouse anti-b-actin (1:30,000), rabbit anti-GluK2/3 (1:2000), and mouse anti-postsynaptic density 95 (PSD95) (1:5000).

Blots were then washed 3 x 5 minutes in TBS and incubated for 1 hour in blocking solution containing secondary antibodies: IRDye680RD donkey anti-rabbit IgG and IRDye800CW donkey anti-mouse IgG diluted (1:10,000), washed 3 x 5 minutes in TBS, rinsed in deionized water and visualized on a Li-Cor Odyssey Fc fluorescent imager.

### 2.4. Immunohistochemistry (IHC)

Mice were anesthetized with ketamine (75 mg/kg) and xylaxine (10 mg/kg) in 0.9% saline before decapitation. Brains were frozen slowly in isopentane before embedding in OCT compound and storage at −80°C. Coronal sections (50 μm) were prepared on a cryostat at −20°C and mounted on glass slides. Sections were thawed to room temperature before carrying out three washes with phosphate buffered saline (PBS), fixed in Carnoy solution (1:6 acetic acid : ethanol) for 20 min, and washed once with PBS. In a humid chamber, the sections were permeabilized in 0.1% Triton X-100 in PBS for 30 min and blocked with 0.05% Tween and 5% newborn calf serum in PBS for another 30 min. After cooling to 4°C, they were incubated in rabbit anti-GluK2/3 antibody (Invitrogen, PA1-37780; 1:100) overnight, washed with PBS three times, and then incubated with goat anti-rabbit-488 (Abcam, ab150077; 1:200) for 2 h at room temperature in the dark. They then were washed with PBS three times and dried for 5 min. Antifade Mounting Medium with DAPI (Vectashield, H-1500-10) was added, and the coverslips were left to cure overnight at room temperature. Images were obtained using a Fluoview FV10i confocal laser scanning microscope (Olympus Life Science, Waltham, MA, USA).

### 2.5. Kainic acid-induced seizures

WT and mutant mice were weighed and injected intraperitoneally (i.p.) with either 5 mg/kg or 25 mg/kg of kainic acid (KA) in a volume of 10 ml/kg. They were then immediately placed in a chamber and video recorded for up to 120 min. Seizure severity was assessed using a modified Racine scale (RS): 1, behavioral arrest; 2, forelimb clonus and/or Straub tail and facial automatisms; 3, repetitive scratching, circling, and/or forelimb clonus with loss of balance; 4, forelimb clonus with rearing and/or falling; 5, repetition of stage 4 behaviors; 6, generalized tonic-clonic seizures, wild running, and/or jumping; 7, death (Racine, 1972). All mice were euthanized after 120 min.

### 2.6. Handling-induced seizures

Handling-induced seizures were measured in T660K and WT littermates once per week beginning in the third postnatal week for 8 weeks and then every other week until the mice were 14 weeks old as described previously (Hawkins et al., 2021). For testing, each mouse was removed from its home cage, placed in a clean cage, and scored for 1 min for the presence or absence of a seizure by an experimenter blinded to genotype. Behavioral seizures in T660K mice ranged in severity and included rearing and forelimb clonus, wild running and jumping, and tonic-clonic seizures. WT mice did not exhibit any seizure behaviors.

### 2.7. Video electroencephalograms (Video-EEG)

EEG headmounts (Pinnacle Technology, Lawrence, KA, USA) were implanted on 2- to 3-month-old male and female WT, A657T, and T660K mice while anesthetized with ketamine/xylazine and head-fixed in a stereotactic frame (Hawkins et al., 2021). Four stainless steel screws in the headmounts acted as cortical surface electrodes and were located bilaterally 1 mm lateral from the midline and 0.5-1 mm anterior to bregma or 4.5-5 mm posterior to bregma. Recordings from the posterior screws were the EEG1 channel and are shown in the figure and used for analysis. The left anterior screw was the ground. Mice were allowed to recover for 2 days after surgery before initiating the video-EEG in a recording chamber. EEG data was collected continuously via a tether attached to the headmount while video captured mouse behaviors using Sirenia acquisition software (Pinnacle Technology) for at least 3 days. Spiking was analyzed using the Sirenia Seizure application and were included in the dataset if they had spike and slow wave morphology and amplitudes of at least 3 times the root mean squared baseline noise.

### 2.8. Open Field test and naturalistic behavior quantification

Open field testing (OFT) was performed on mice separated by sex. Each group was brought to the testing room, allowed to acclimate for 30 min, and then placed in the center of a 60 cm x 60 cm x 31 cm acrylic arena for 10 min. Spontaneous digging behavior was assessed by placing mice into a behavioral chamber filled with bedding for 10 min. Ethovision (Noldus, Leesburg, VA, USA) was used to track and quantify locomotion from open field (OF) recordings. Naturalistic behaviors including grooming, rearing, and digging were manually quantified by a blind experimenter using the free and open-source event-logging software BORIS (Friard and Gamba, 2016). The time spent in the center 36.5 cm x 36.5 cm square of the box was quantified using BORIS.

### 2.9. Zero Maze test

Mice were placed on an elevated zero maze for 10 min while their movement was recorded using Ethovision or LimeLight motion tracking software (Actimetrics, Lafayette, IN, USA) in the Northwestern University Behavioral Phenotyping Core. The maze consisted of a circular track (56 cm diameter) divided into four quadrants with alternating open and closed walls. The percentage of time that animals spent in the closed quadrants over a total of 10 min was measured with LimeLight or Ethovision.

### 2.10. Nestlet shredding

Animals were individually housed and given a pre-weighed piece of cotton nestlet overnight (∼2.5 g). The remaining unshredded Nestlet was removed the following morning and allowed to dry overnight. It was then weighed to determine the percent of the Nestlet shredded (Angoa-Perez et al., 2013).

### 2.11. Rotarod test

Mice were trained and tested on a rotarod over a two-day period for three trials per day. On day one, animals were trained to stay on the rotarod test while facing forward. On the second day, the mice were tested on three trials; times spent on the rod during second and third trials were averaged to quantify performance. Before each trial, mice were placed onto the rotarod at a starting speed of 4 rpm. The rotarod accelerated to 40 rpm over the course of 180 sec and the latency to fall was recorded for each mouse. Mice that did not fall during the testing period were recorded as 180 sec. During training, mice that fell off within the first 20 sec were placed back onto the rod. Testing trials where mice did not face forward were excluded.

### 2.12. Four-limb Hang

Mice were acclimated to the testing room for at least 30 min. Each mouse was placed on a wire cage lid, gripping with all four limbs. The lid was then inverted 30 cm above a padded plastic container, suspending the mice for up to 6 min. The time until each animal fell was recorded.

### 2.13. Balance beam test

Traversal across a balance beam was performed as described previously (Luong et al., 2011). The balance beam consisted of a 1 m long polyurethane-sealed wooden beam with a 6 mm surface suspended 40 cm above the tabletop and held at a 2° angle. A safe box was placed at the end of the balance beam and held nesting material from their home cage. The first and last 10 cm of the beam were used as the starting and ending zones and traversal in only the center 80 cm was analyzed. Mice were trained for one day and tested the next day. Each group was separated by sex and given 30 min to habituate to the room prior to testing. Mice were placed before the starting line and were trained to cross the beam without stalling. After each trial, mice were allowed to rest in the safe box for 15 s before returning to their cage for 10 min before their next trial. Mice performed three trials each day. On the testing day, the first trial was used as a practice trial, and the second and third trials were averaged as to score performance. Trials where mice stalled were omitted, and an additional trial was performed as a replacement. Mice that did not successfully complete two testing trials were omitted. Latency to cross and number of paw slips were scored by a blinded observer using BORIS (Friard and Gamba, 2016). 3 WT animals, 1 A657T, and 1 T660K animal did not run for a consecutive meter across the balance beam twice and were not included in analysis.

### 2.14. Gait analysis

The DigiGait (Mouse Specifics Inc., Framingham, MA, USA) rodent treadmill system in the Northwestern University Behavioral Phenotyping Core was used to quantify various aspects of GluK2(A657T) gait. The parameters analyzed were limited to those that were consistently different during preliminary testing to minimize multiple comparison effects. Mice were first habituated to the treadmill running at 24 cm/s for about 2 s and were then tested at the same speed. Videos ranging from 3-6 sec including at least 10 steps of continuous gait for each limb were obtained. 2 GluK2(T660K) mice had repeated seizures and were unable to complete the testing. DigiGait software was unable to effectively track limbs for 2 WT (T660K line) and 2 T660K. These mice were not included in the analysis. Results are presented as averages of both the right and left hindlimbs or forelimbs.

### 2.15. Ataxia phenotyping

Mice were rated in ledge navigation, hindlimb clasping, and kyphosis tests on the Guyenet ataxia phenotype scoring system (Guyenet et al., 2010). Performance and posture in each test were scored on a scale of 0-3, with 0 indicating that the phenotype is not present and a score of 3 given to those animals with the most severe phenotype, by 2 blind experimenters and then averaged. Average scores from all 3 tests were averaged to obtain a composite ataxia score. For the ledge test, animals were placed on the edge of an empty cage. Mice typically remained still, turned circles, walked along the ledge, and/or lowered themselves into the cage.

Animals that descended into the cage or walked along the ledge without foot slips were given a score of 0. Animals that incurred foot slips were given a score of 1. Animals that could not effectively use their hindlimbs were scored 2. Animals that nearly fell, fell, or refused to move were scored 3. For the kyphosis test, animals were placed on an open, flat table and allowed to traverse freely. Mice were scored based on the presence of spine curvature that persists during locomotion and were scored as: 0: no kyphosis, 1: mild kyphosis with ability to straighten spine, 2: persistent mild kyphosis, and 3: persistent severe kyphosis. Hindlimb clasping was scored during 10 s of tail suspension. Scoring was as follows: 0: hindlimbs were consistently splayed outward, 1: one hindlimb retracted the majority of the time, 2: both hindlimbs retracted the majority of the time, 3: both hindlimbs are entirely retracted the majority of the time.

### 2.16. Tail suspension test

Animals were suspended by their tails to a flat ledge using tape and allowed to hang for six min. A 4 cm long piece of tubing was placed over their tails to prevent climbing. Percent time immobile was calculated from 6 min of video using BORIS. Immobility of less than 2 s in duration was removed from analysis.

### 2.17. Climbing scoring

Mice were scored on their ability to right themselves to grasp their tails during 6 min of tail suspension with a system adapted from (Hawkins et al., 2024). Animals were scored: 1: able to curl/climb up tail to right self, 2: attempts to curl up and right self unsuccessfully, 3: no attempt to curl up.

### 2.18. Dystonia scoring

Mice were scored for presenting dystonic-like behaviors during 2 min of tail suspension adapted from a similar scale in (Su et al., 2023). Dystonic-like behaviors included fore/hind-paw clenching, hindlimb clasping, and sustained hindlimb hyperextension. Scores ranged from 0-3 in ascending severity: 0: no dystonic-like behaviors, 1: dystonic-like behaviors <50% of the duration, 2: dystonic-like behaviors >50% of the duration, 3: dystonic-like behaviors 100% of the duration.

### 2.19. T-maze test

Automated T-Maze testing (MazeEngineers, Skokie, IL, USA) was performed to assess learning and working memory as described (Moya et al., 2023). GluK2(A657T) mice and their WT littermates were singly housed and food restricted to maintain ∼85% of their ad libitum body weight during testing. GluK2(T660K) animals incurred adverse reactions including lethargy and death at this level of food restriction, so this line and their WT littermates were food restricted for only 2 hours prior to testing. The mice underwent two days of habituation where they were able to explore the maze for 30 min while receiving food reward for their exploration. On day 1, mice were incentivized to explore each arm of the maze by receiving chocolate-flavored pellets from automated food dispensers each time they entered a choice arm. On day 2, pellets were only obtained from the food dispensers upon entry to a choice arm. Mice then underwent pre-training where they completed 20 forced-choice trials in alternating directions until they were able to complete the trials in under 30 min for two consecutive days. Animals then began delayed non-match to sample (DNMS) training during which they performed 10 sets of a forced trial paired with subsequent choice trial. Mice were delayed for 2s between the forced and choice trials. Animals were rewarded when they successfully chose the alternate arm from the forced trial. Mice continued with DNMS training until they reached ≥ 70% correct choices for three consecutive days. After training, animals moved with delays of 2s, 10s, and 30s prior to the choice trial on two consecutive days. One WT animal (an A657T littermate) and 4 T660K mice that did not reach ≥ 70% performance by day 8 were excluded from the study. Days to criteria were recorded and compared between genotypes for animals that met criteria. T-Maze performance was averaged for each delay and compared between *Grik2* mutants and their WT littermates.

### 2.20. Novel object recognition

Habituation for novel object recognition (NOR) testing was performed in a 61 cm x 61 cm x 30 cm OF chamber. One day after habituation, animals were separated by sex and allowed 30 min to acclimate to the room prior to completing training and novel object recognition testing. The mice were given a 1 hr inter-trial interval (ITI) between training and testing. During testing, animals were placed in the arena for 10 min and allowed to explore the arena, which had 4 quadrants in which to place objects. The objects were placed in adjacent quadrants during training. After training, one of the two objects was moved diagonally to the distal quadrant from the familiar object for a subsequent training period. For NOR testing, the object in the novel location was replaced by a novel object. Objects were placed at an equal distance from each other and the sides of the arena (Denninger et al., 2018). A glass beaker filled with teal sand and a flask filled with marbles were selected as the object to be used during training. A colorful jar was used as the novel object during NOR. A preliminary cohort of mice that was given 5 minutes to explore did not show bias towards any of these objects. Animals were grouped based on sex and genotype and pseudo-randomly assigned to an object that would be removed and replaced for NOR. Exploration was scored when animals were engaged and facing the object within a 2 cm radius of the object using BORIS software. Time spent grooming, climbing, and orienting away from the object were excluded. The discrimination index between the familiar and novel objects was calculated as (novel object exploration time - familiar object exploration time)/(novel object exploration time + familiar object exploration time) (Denninger et al., 2018).

### 2.21. Developmental testing

Neonatal development was assessed with motor tests and ultrasonic vocalization recordings at postnatal days 3, 7, and 10 (P3, P7, and P10) in the Northwestern University Behavioral Phenotyping Core. Pups were weighed on each day of testing. To test ambulation, each pup was placed into a standard rat cage (46 cm x 25 cm x 23 cm) and allowed to move freely for 3 min while video was recorded from above and from the side of the cage. Animals were scored on an ordinal scale from 0-3 with 0: no movement, 1: crawling with asymmetric limb movement, 2: slow crawling with symmetric limb movement, and 3: fast crawling/walking (Feather-Schussler and Ferguson, 2016). Righting reflexes were measured after inverting pups to their backs on a flat surface and recording the time to return to a prone position. Three trials per pup were performed. Trials were stopped after one minute if righting had not occurred. Hindlimb strength and neuromuscular function were assessed using hindlimb suspension tests. Pups were hung head down from their hind limbs on the rim of a 50 ml conical tube for up to one minute per trial for 3 trials. The latency to fall was measured and hindlimb posture of each animal was scored from 0 to 4: 4: normal separation, 3: less separation with infrequent touching, 2: minimal separation and frequent touching between hindlimbs, 1: clasping the majority of the time, and 0: constant clasping and/or fall (Feather-Schussler and Ferguson, 2016).

### 2.22. Ultrasonic vocalizations

Ultrasonic vocalization (USV) recordings were obtained from pups using the Pettersson Ultrasound Detector D500X (Bat Conservation and Management, Inc., Carlisle, PA, USA) inside a soundproof box supplied by the Northwestern University Behavioral Phenotyping Core. An individual pup was placed in the chamber that was covered with a lid containing a microphone and recorded for 5 min. USVs were identified and analyzed using MUPET software (Van Segbroeck et al., 2017).

### 2.23. Quantification and statistical analysis

Statistical analyses were performed using GraphPad Prism (GraphPad Software, Boston, MA). Unpaired comparisons were made using either the Student’s t-test for normally distributed data or the Mann-Whitney test. Normality was determined using the Shapiro-Wilk test. Multiple comparisons were performed using two-way ANOVA followed by post hoc Sidak’s multiple comparisons test to compare rows or Tukey’s if row and column comparisons were required. Differences were considered significant if p<0.05.

## 3. Results

We generated mouse models for the two most common human pathological variants in the *GRIK2* KAR gene, p.Ala657Thr and p.Thr660Lys. Individuals with these *de novo* variants have NDDs with ID and GDD as prominent features. Earlier biophysical studies of recombinant GluK2(A657T) and GluK2(T660K) receptors suggested that these variants confer a GoF on KARs containing the mutant subunits (Guzmán et al., 2017; Stolz et al., 2021). A focused neurophysiological analysis of hippocampal function in the A657T mice supported the GoF categorization (Nomura et al., 2023). Here we analyze behavioral phenotypes of the GluK2(A657T) and GluK2(T660K) mice (also denoted A657T and T660K, respectively) to determine the extent to which they recapitulate the dysfunction in individuals with each variant.

### 3.1. Grik2 GoF variants reduce viability

The A657T and T660K mouse lines were generated by introducing orthologous variants in *Grik2* - c.1969G>A, p.Ala657Thr or c.1979C>A, p.Thr660Lys, respectively - on a C57BL/6J background using CRISPR-Cas9 editing (Fig. 1A). Both lines of mice are fertile as heterozygotes, but dams display reduced viability during parturition, so lines were maintained with male breeders. Breeding in which both parents were heterozygous yielded no living homozygous pups (A657T, n=80; T660K, n= 31 pups analyzed), consistent with embryonic lethality. Neither line generated pups at Mendelian ratios in heterozygous crosses with WT C57Bl/6 mice: A657T offspring were born at 75% the rate of WTs (61 A657T:79 WT mice), while T660K mice were born at a marked 33% of the WT rate (59 T660K:179 WT mice). All mice used for experiments in this study therefore were heterozygous A657T, heterozygous T660K, or each of their respective WT littermates. Both A657T and T660K animals display significantly reduced body mass than their WT littermates (A657T males: F_1, 16_ = 33.2, P<0.0001, A657T females: F_1, 18_ = 11.4, P=0.0033; T660K males: F_1, 22_ = 59.3, P<0.0001, T660K females: F_1, 21_ = 31.9, P<0.0001; Fig. 1B). GluK2(T660K) mice of both sexes were already smaller than their WT littermates as early as 3 weeks of age (males: T660K: 8.6 ± 1.7 g, n = 9; WT: 11 ± 1.6 g, n = 15, p=0.014; females: T660K: 8.7 ± 1.1 g, n = 8; WT: 10 ± 1.2 g, n = 15, p=0.011), while A657T were significantly smaller than WTs beginning at 5 weeks for males and 7 weeks for females (males: A657T: 17 ± 0.93 g, n = 8; WT: 19 ± 0.92 g, n = 10, p=0.0038; females: A657T: 16 ± 0.76 g, n = 8; WT: 18 ± 1.2 g, n = 12, p=0.041; Fig. 1B).

**Figure 1.**
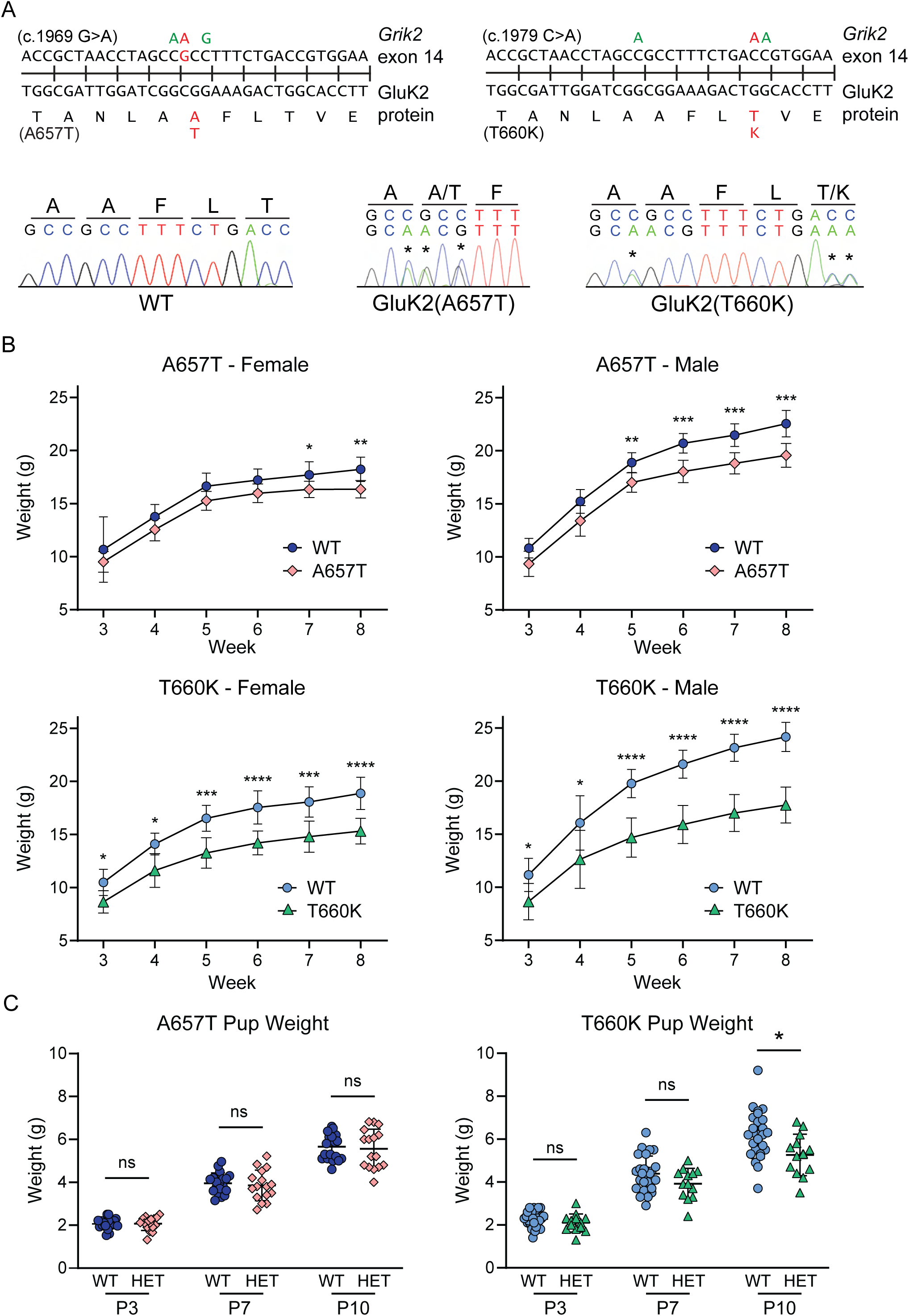
Genetic background and growth curves of GluK2(A657T) and GluK2(T660K) mutant mice. A) Top: Nonsynonymous point mutations c.1969G>A and c.1979C>A are shown in red in the mouse *Grik2* gene. The consequent p.Ala657Thr (A657T) and p.Thr660Lys (T660K) substitutions are analogous to pathogenic variants in humans. Silent mutations are shown in green. Bottom: Chromatograms from Sanger sequencing of the knockin mice. Asterisks mark the heterozygous alleles edited in the knockins. B) Average weights ± SEMs of WT, A657T, and T660K animals from 3 to 8 weeks of age. A657T and T660K mice of both sexes weighed less than age-matched WT littermates. A657T females and A657T males began to diverge from their WT littermates at weeks 7 and 5, respectively. T660K mice of both sexes weighed less than WT mice when tested after weaning in week 3. C) Pup weights were recorded to evaluate early postnatal growth and viability. A657T and their WT littermates weighed the same, but T660K mice are smaller starting at P10. Asterisks indicate p-values: *, p < 0.05; **, p < 0.01; ***, p < 0.001; ****, p < 0.0001.

To determine if *Grik2* GoF display early deficits in growth, we weighed them at three early developmental timepoints – P3, P7, and P10. From P3 to P10, A657T pups weighed the same as their WT littermates (genotype effect: F_1, 33_ = 0.135, P=0.72), indicating that they thrive to the same degree (Fig. 1C). These data are also consistent with the later-developing divergence in WT and A657T mice weights (Fig. 1B). GluK2(T660K) mice, in contrast, weigh significantly less than their WT littermates starting at P10 (genotype effect: F_1, 37_ = 4.73, P=0.036; P3: T660K: 2.1 ± 0.42 g, n = 13; WT: 2.3 ± 0.37 g, n = 26, p= 0.50; P7: T660K: 3.9 ± 0.71 g, n = 13; WT: 4.4 ± 0.82 g, n = 26, p= 0.22; P10: T660K: 5.3 ± 0.97 g, n = 13; WT: 6.2 ± 1.1 g, n = 26, p=0.034; Fig. 1C), indicating that this line may suffer from early reductions in viability beyond embryonic lethality.

### 3.2. Grik2 variants alter GluK2-containing KAR function, expression, and localization

The *GRIK2* variants exhibit putative GoF characteristics in recombinant systems, but they appear to have differing effects on expression and localization in mice. In analogous recombinant mutant receptors, both variants exhibit greatly increased affinity for glutamate, a destabilized closed state of the channel, and prolonged current decay (Guzmán et al., 2017; Kohda et al., 2000; Stolz et al., 2021). As well, KAR excitatory postsynaptic currents (EPSC_KA_) in hippocampal CA3 pyramidal neurons from GluK2(A657T) mice exhibited a much slower decay time course compared to WT and gated a tonic current in response to ambient glutamate, consistent with the GoF channel properties in recombinant analyses (Nomura et al., 2023). In A657T mice, expression of the GluK2/3 protein appeared normal in cortical, striatal, hippocampal, and cerebellar regions (Fig. 2C) (Nomura et al., 2023). In contrast, GluK2/3 staining intensity appeared diminished and differently localized in IHC images in hippocampal regions in GluK2(T660K) mice, which lack the striking enrichment of GluK2/3 staining in *stratum lucidum* arising from the predominant localization of KARs to mossy fiber synapses (Fig. 2C). These observations suggest that the mutant T660K subunit may have a dominant-negative effect on KAR expression and synaptic targeting, which is supported by Western blots showing GluK2/3 immunoreactivity is significantly reduced in T660K hippocampal P3 fractions (Fig. 2A, B). In contrast, GluK2/3 expression in P3 fractions was not previously found to be different in A657T animals (Nomura et al., 2023).

**Figure 2.**
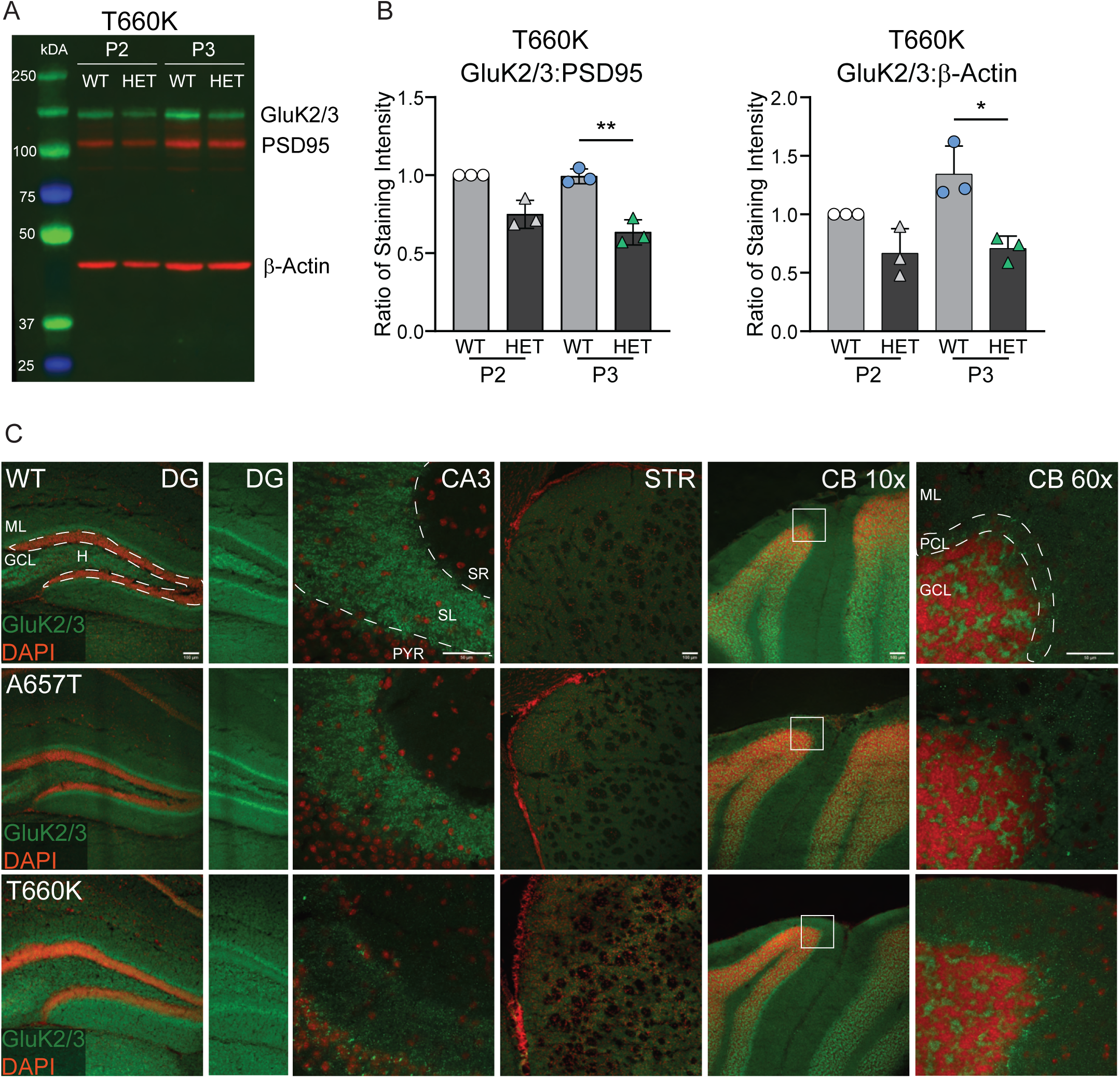
GluK2/3 protein expression and localization are grossly normal in A657T mice but aberrant in T660K mice. A) Representative Western blot of GluK2/3, PSD-95, and b-actin immunoreactivity in WT and T660K hippocampal P2 (small synapses) and P3 (mossy fiber bouton synapses) fractions. B) Quantification of Western blots with band densities normalized to WT P2 bands within each experiment. The ratio of GluK2/3 to both PSD-95 and b-actin was reduced in T660K P3 fractions. An equivalent Western blot analysis showed GluK2/3 expression was unchanged in A657T mice (Nomura et al., 2023). Asterisks indicate p-values: * p < 0.05, ** p < 0.01. C) Representative confocal images of GluK2/3 expression and DAPI counterstain in coronal sections of the hippocampal dentate gyrus (DG), CA3 region (CA3), striatum (STR), cerebellar vermis and simple lobule (CB 10x), and a further 6x zoomed image of the tip of the simple lobule (CB 60x). Gross morphology was not obviously altered between genotypes. DG: GluK2/3 labeling is observed throughout the molecular layer (ML) and hilus (H) and is enriched in the inner molecular layer. GluK2/3 labelling is present in the IML of T660K animals but is also prominent in the granule cell layer (GCL). CA3: WT and A657T mice show characteristic enrichment in the *stratum lucidum* (SL) compared to the *stratum radiatum* (SR). T660K mice, in contrast, show reduced expression in the SL, and bright puncta in the pyramidal cell layer (PYR). STR: Striatal expression of GluK2/3 is generally diffuse and not notably different between genotypes. CB 10x and CB 60x: Cerebellar localization of GluK2/3 is not notably different between genotypes and is primarily observed in the granule cell layer (GCL) with diffuse staining in the Purkinje cell (PCL) and molecular (ML) layers. Scale bars: DG: 100μm, CA3: 50 μm, STR: 100 μm, CB: 100 μm, CB 6x: 50 μm.

### 3.3. Grik2 GoF mice are acutely sensitive to KA-induced seizures

To resolve whether the A657T and T660K mutations cause a GoF or LoF in KAR signaling, we tested the sensitivity of the mice to seizures induced by the eponymous agonist KA, which has been used for decades to evoke acute and chronic hippocampal-dependent seizures as a model of temporal lobe epilepsy (TLE) (Ben-Ari, 1985). Initiation of seizures by KA is dependent upon activation of hippocampal GluK2-containing KARs (Mulle et al., 1998). We therefore predicted that the mutant mice would have heightened sensitivity to KA-induced seizures if they indeed have a GoF in KAR signaling.

To test this prediction, we injected A657T and T660K mice with a standard dose of 25 mg/kg KA and measured seizure severity as the time to reach a modified Racine score of 4 (RS4, forelimb clonus with rearing and/or falling; see Methods) (Fig. 3A). Both mutant lines reached RS4 significantly faster than their WT littermates (A657T: 5.3 ± 1.8 min, n = 9; WT: 38 ± 16 min, n = 11, p<0.0001; T660K: 5.9 ± 2.8 min n = 12; WT: 37 ± 16 min, n = 10, p<0.0001; Fig. 3A). Moreover, all mutant mice rapidly progressed in seizure severity to death (A657T: 11 ± 2.3 min; T660K: 14 ± 7.3 min), whereas only 2 animals in each WT group died before termination of the experiments at 120 min.

**Figure 3.**
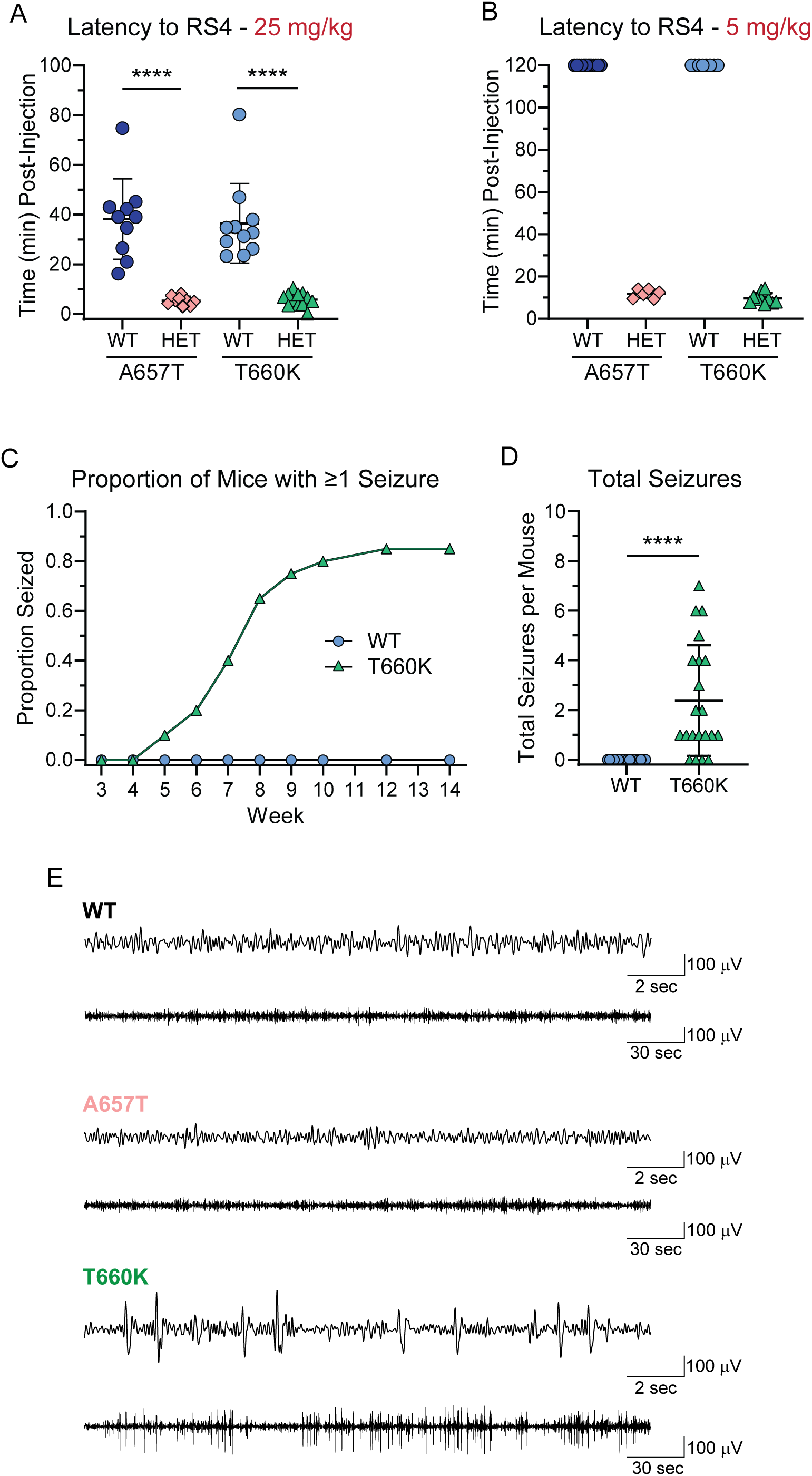
A657T and T660K mice are more susceptible to KA-induced seizures, and T660K mice display aberrant EEGs and handling-induced seizures. A) Latency to reach RS4 after 25 mg/kg KA injection is reduced for both A657T and T660K mice. B) A657T and T660K mice were acutely sensitive to a low dose of KA (5 mg/kg) that does not cause convulsant or subconvulsant behaviors in WT animals. The latency to reach RS4 was ∼10 min for the mutant mice. C) Handling induced seizures were recorded once per week for 10 weeks beginning at the third week postnatal. WT mice did not display behavioral seizures, while 85% of T660K mice exhibited at least one seizure during the testing period. D) T660K mice exhibited an average of 2.4 seizures during the 10-week testing period, while WTs exhibited none. E) Cortical scalp EEG recordings from WT, A657T, and T660K mice are shown at two timescales. Interictal cortical hyperactivity was only observed in T660K mice. Asterisks indicate p-values: ****, p < 0.0001.

We then reduced the dose of KA to 5 mg/kg to test if the enhanced seizure severity observed in the mutant mice was accompanied by an increased potency of the chemoconvulsant. WT mice did not have seizures or exhibit subconvulsant behaviors at this dose of KA during the post-injection observation period of 120 minutes. In contrast, all mutant mice seized, reaching RS4 at 11.8 ± 2.1 min (A657T, n = 6) or 9.7 ± 2.5 min (T660K, n = 10; Fig. 3B). All A657T mice died during the test, whereas half of the T660K survived to 120 minutes despite having continuous seizures of varying severity. These data demonstrate that both A657T and T660K mutations greatly enhance sensitivity of the mice to induction of behavioral seizures and lethality by KA and therefore indeed are GoF.

### 3.4. GluK2(T660K) mice exhibit abnormal EEGs and handling-induced seizures

We noted early in our analysis that T660K mice often exhibited spontaneous seizures during routine handling, which was not observed in either A657T or WT mice. Similarly, individuals with the *GRIK2* p.Thr660Lys variant, but not the p.Ala657Thr variant, suffer from epilepsy. We therefore measured the occurrence of clonic or clonic-tonic seizures in T660K mice during handling once a week between 3 to 14 weeks of age. No WT animals displayed seizures, while T660K mice began seizing as early as 5 weeks of age (Fig. 3C). By 14 weeks of age, 85% of T660K animals had seized at least once and in total incurred an average of 2.4 seizures over the testing period (Fig. 3C, D).

We carried out video-EEGs to determine if any of the mice showed cortical hyperexcitability in the absence of behavioral seizures. Electrographic recordings from two pairs of cortical scalp electrodes were obtained for at least 72 hrs from each of the strains of mice. Neither WT nor A657T mice exhibited abnormal spike activity, whereas five of six T660K mice exhibited long trains of low-frequency (<1 Hz) spike and wave complexes interspersed with delta and higher frequency bursts in both EEG channels in the absence of subconvulsant behaviors, as shown in the representative examples at two time bases in Fig. 3E. These data therefore demonstrate that the T660K mice display frequent cortical hyperactivity in interictal states, consistent with their propensity for spontaneous behavioral seizures.

### 3.5. Grik2 GoF mice exhibit altered activity and a reduction in naturalistic behaviors

Individuals with the *GRIK2* p.Ala657Thr GoF variant have proprioceptive dysfunction and unstable gaits, whereas children with the severely debilitating p.Thr660Lys variant are not ambulatory and show limited responsiveness to external stimuli (Guzmán et al., 2017; Stolz et al., 2021). We therefore tested if the mouse models of these disorders exhibited similar divergence in locomotion, naturalistic, or other behaviors that are compromised in many NDD mouse models. We started with an open field movement assay (Fig. 4A). When comparing the total distance traveled in 10 min, A657T mice were equivalent to that of WT littermates, whereas T660K mice moved further than WTs (A657T: 4000 ± 1200 cm, n = 15; WT: 4300 ± 870 cm, n = 15, p=0.39; T660K: 5300 ± 1800 cm, n = 22; WT: 4400 ± 970 cm, n = 30, p=0.022; Fig. 4A, B). When the distance traveled was binned by minute, however, it became evident that both A657T and T660K mice moved significantly less than their WT littermates during the first minute of OFT (A657T: 200 ± 120 cm, n = 15; WT: 600 ± 200 cm, n = 19, p<0.0001; T660K: 200 ± 140 cm, n = 22; WT: 630 ± 280 cm, n = 30, p<0.0001; Fig. 4D). Locomotor activity in the two lines diverged subsequently, with the A657T mice normalizing to WT distance traveled, while the T660K mice were hyper-locomotive during the second half of the testing period (minute 6: T660K: 580 ± 190 cm, n = 22; WT: 400 ± 150 cm, n = 30, p=0.0080; Fig. 4E). These results demonstrate that A657T and T660K mice displayed aberrant locomotor activity characterized by initial limited mobility followed by normalization (A657T) or hyperactivity (T660K).

**Figure 4.**
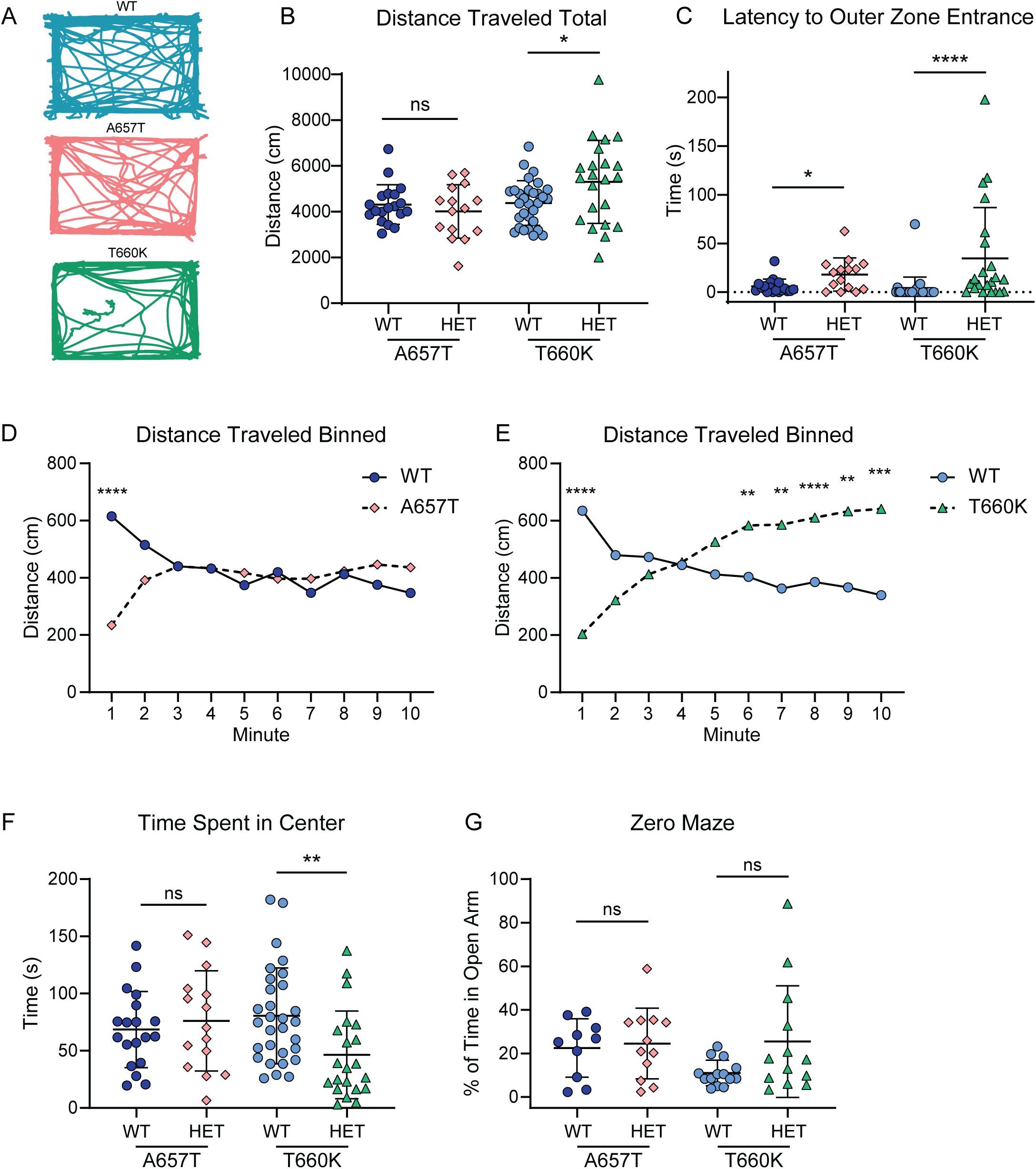
*Grik2* GoF mice exhibit altered activity in open field testing. A) Representative movement traces from 10 min of OF exploration. B) Total distance traveled during 10 min of OF exploration was the same for A657T and WT mice, but T660K mice traveled farther than WTs. C) The latency to enter the outer zone was measured after placing the mice in the center of the OF chamber. A657T and T660K mice took longer than WTs to move to the outer zone. D) Total distance traveled was binned by minute to assess locomotor activity over the 10 min. A657T mice exhibited reduced movement compared to wild-type controls during the first minute, but activity normalized thereafter. E) Total distance traveled binned by minute shows that T660K mice also moved less than WTs during the first minute but were hyperactive in the last half of the test. F) The time spent in center was measured during OFT to evaluate anxiety-like behavior. A657T mice spent the same amount of time as WTs, while T660K mice spent less time in the center. G) The percent of total time spent in the open arms of the zero maze was compared between genotypes to assess anxiety-like behavior. *Grik2* GoF mice spent the same amount of time in open arms as WTs. Asterisks indicate p-values: *, p < 0.05; **, p < 0.01; ***, p < 0.001; ****, p < 0.0001.

We assessed if anxiety might contribute to the limited movement during the first minute of OFT by comparing the time spent in the center of the OF chamber with the time spent in the outer region. When mice were initially placed in the center of the OF, A657T and T660K animals exhibited a longer latency to their first transition from the center to the outer zone than their WT littermates (A657T: 18 ± 17 s, n = 15; WT: 6.0 ± 7.4 s, n = 19, p=0.044; T660K: 35 ± 52 s, n = 22; WT: 2.9 ± 13 s, n = 30, p<0.0001; Fig. 4C). The initial immobility and increased latency to transition could be interpreted as freezing. When averaged across all 10 minutes, however, A657T mice spent the same amount of time in the center of the OF as their WT littermates (A657T: 76 ± 44 s, n = 15; WT: 69 ± 33 s, n = 19, p=0.57; Fig. 4F). T660K animals, in contrast, spent significantly less time exposed in the center of the OF than their WT littermates, as seen in the example traces (T660K: 46 ± 38 s, n = 21; WT: 81 ± 42 s, n = 30, p=0.0017; Fig. 4A, F).

Freezing in mice is a behavior typically attributed to anxiety or fear. To further explore the relevance of anxiety to the *Grik2* locomotor phenotypes, we performed a zero-maze test in which mice traverse a circular track with opposing open and enclosed quadrants. Both lines of mutant mice spent an equivalent percent of time in the open regions of the maze as their WT counterparts (A657T: 25 ± 16 %, n = 12; WT: 23 ± 13 %, n = 10, p=0.75; T660K: 25 ± 26 %, n = 13; WT: 11 ± 5.9 %, n = 14, p=0.15; Fig. 4G), suggesting that these mutants do not display generalized anxiety-like behavior.

We next examined if naturalistic behaviors associated with perseverative phenotypes were altered in the *Grik2* GoF mice. Rearing and grooming behaviors in the OF were compared between the WT and *Grik2* mutant mice (Fig. 5A-B). We observed that T660K mice groomed significantly less than their WT littermates (T660K: 6 ± 7 s, n = 12; WT: 22 ± 10 s, n = 14, p=0.0002), while A657T mice groomed an equivalent amount (A657T: 23 ± 13 s, n = 16; WT: 27 ± 10 s, n = 15, p=0.55; Fig. 5A). The *Grik2* mutant mice reared significantly less often than their WT littermates (A657T: 38 ± 23 s, n = 16; WT: 67 ± 13 s, n = 15, p= 0.0004; T660K: 40 ± 24 s, n = 11; WT: 62 ± 28 s, n = 14, p=0.050; Fig. 5B). Surprisingly, both *Grik2* mutant lines failed to dig when placed in an OF chamber with bedding (Fig. 5C). In contrast, WT mice spent over one minute digging on average over the 10 minutes of testing (A657T: 0.50 ± 0.53 s, n = 10; WT: 140 ± 130 s, n = 10, p<0.0001; T660K: 2.4 ± 3.4 s, n = 17; WT: 83 ± 39 s, n = 22, p<0.0001; 5C). To complete our analysis of naturalistic behaviors, we compared the extent to which mice shredded nestlets provided for building nests in home cages. Both *Grik2* mutant mice displayed a reduction in nestlet shredding compared to their WT littermates (A657T: 66 ± 38 %, n = 12; WT: 93 ± 8.9 %, n = 10, p=0.04; T660K: 23 ± 35 %, n = 17; WT: 88 ± 20 %, n = 22, p<0.0001; Fig. 5D), consistent with reductions in other naturalistic behaviors. The behavioral test battery therefore shows that *Grik2* mutant mice perform naturalistic behaviors with reduced frequency and do not display perseverative behaviors relative to their WT counterparts.

**Figure 5.**
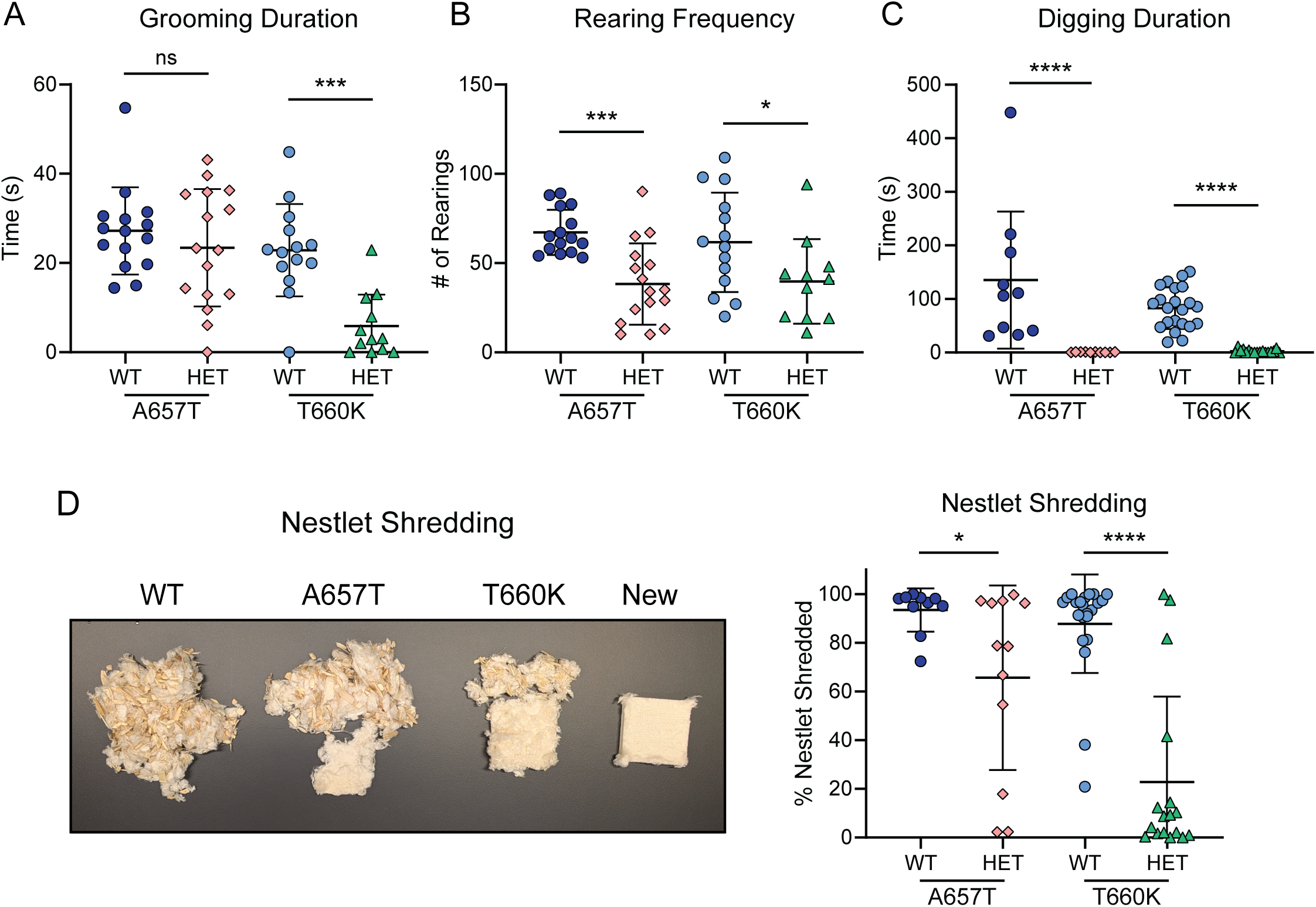
A657T and T660K mice perform naturalistic behaviors at reduced rates. A) A657T mice groomed as much as WTs, but T660K mice groomed less during the 10 min test period in an OF chamber. B) Both A657T and T660K animals reared less than their WT littermates during 10 min of OFT. C) *Grik2* GoF mice failed to dig in woodchip bedding when observed in an OF over 10 min. WT digging was normal. D) Representative images of nestlets shredded by mice during nest-building behaviors compared to a new nestlet. The percentage of each nestlet shred during an overnight period was lower for the A657T and T660K mice. Asterisks indicate p-values as follows: *, p < 0.05; ***, p < 0.001; ****, p < 0.0001.

### 3.6. Grik2 mutant mice display impaired balance and altered gait

Motor deficits are a recurring feature of human *Grik2* disorders. All individuals with the *GRIK2* p.Ala657Thr variant have gait dysfunction that ranges from ataxia to imbalance, in addition to postural abnormalities and choreiform movements. Children with the p.Thr660Lys variant are not ambulatory and display a variety of symptoms related to abnormal tone including limb hypertonia, truncal hypotonia, spasticity, and opisthotonus. We therefore tested the *Grik2* mutant mice in balance, gait, posture, and motor coordination assays. Mice were tested on an accelerating rotarod to probe for impairments in both motor coordination and balance. A657T mice performed as well as WT littermates (A657T: 99 ± 19 s, n = 13; WT: 82 ± 29 s, n = 15, p=0.08), while T660K mice fell significantly more quickly (T660K: 76 ± 23 s, n = 13; WT: 95 ± 26 s, n = 14, p=0.05; Fig. 6A). Latency to fall in the four-limb hang test was not significantly different between WT and A657T and mutant mice (A657T: 180 ± 80 s, n = 12; WT: 140 ± 77 s, n = 10, p=0. 0.22; Fig. 6B), indicating that these mice do not have compromised limb strength. In contrast, T660K mice fell from the wire lid sooner than WTs during the hang test, suggesting strength deficits may play a role in their motor impairments (T660K: 120 ± 140 s, n = 9; WT: 320 ± 86 s, n = 10, p=0.0066; Fig. 6B). A657T mice crossed an inclined balance beam more slowly (A657T: 5.3 ± 0.69 s, n = 19; WT: 4.3 ± 0.66 s, n = 25, p<0.0001) but did not exhibit more paw slips than their WT littermates (A657T: 2.0 ± 1.9 slips, n = 19; WT: 1.4 ± 1.1 slips, n = 25, p=0.40 Fig. 6C, D), consistent with a mild impairment in balance.

**Figure 6.**
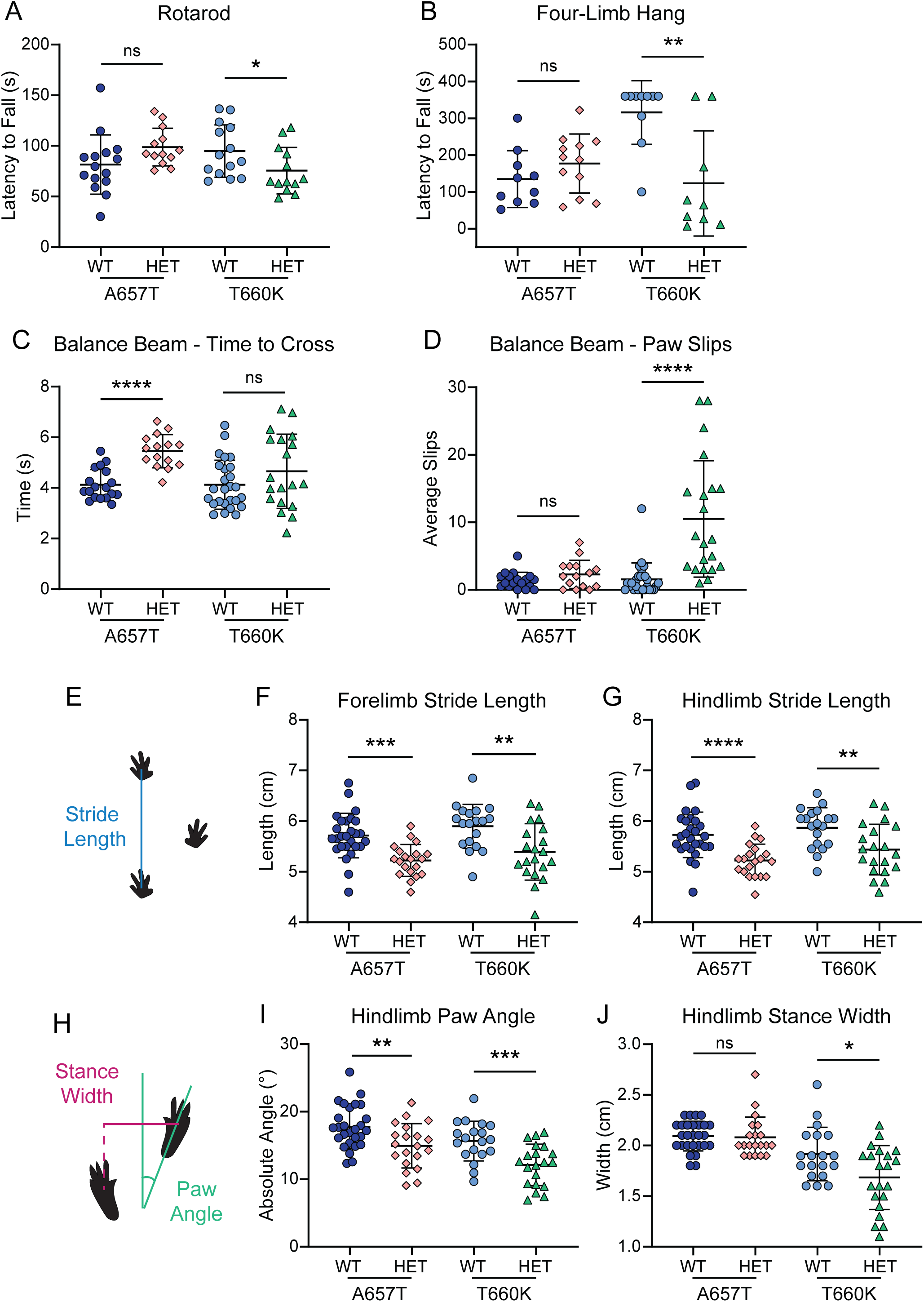
*Grik2* GoF mice exhibit motor deficits in balance and gait. A) Rotarod performance was averaged across 2 trials to assess gross motor coordination and balance. A657T mice performed as well as WTs, while T660K mice performed worse. B) Four-limb hang testing was performed to measure animal limb strength. A657T mice took as long to fall as WTs. T660K mice fell more quickly. C) Latency to cross a balance beam was longer for A657T mice than WTs. T660K mice took the same time to cross as their WT littermates. D) A657T animals did not incur more paw slips during transit across a balance beam. T660K mice slipped more often than WTs. E) Gait was assessed using the DigiGait treadmill apparatus and software. The diagram shows how stride length is measured. F) Forelimb stride length was lower for both A657T and T660K mice. G) Hindlimb stride length was also lower for both *Grik2* GoF mice. H) The diagram demonstrates stance width and paw angle measurements. Stance width is the perpendicular distance between paw centroids during peak stance. Paw angle is measured relative to the axis of the direction of motion. I) Hindlimb paw angle was reduced in both Grik2 GoF mice. J) Hindlimb stance width was reduced in T660K mice, but not in A657T mice. Asterisks indicate p-values: *, p < 0.05; **, p < 0.01; ***, p < 0.001; ****, p < 0.0001.

T660K mice, in contrast, crossed the beam as quickly as their WT littermates (T660K: 4.1 ± 1.0 s, n = 19; WT: 4.7 ± 1.5 s, n = 26, p=0.25), but slipped more frequently (T660K: 11 ± 8.6 slips, n = 21; WT: 1.6 ± 2.4 slips, n = 27, p<0.0001; Fig. 6C, D). The larger number of paw slips but equivalent latency to cross the beam may suggest that the T660K mice were quicker during bouts of movement between paw slips. More generally, these data demonstrate that the *Grik2* mutant mice have balance deficits that manifest in distinct parameters of these tasks.

Gait analysis in the DigiGait treadmill test revealed that both *Grik2* mutant mice ran with a reduced fore- and hindlimb stride length (forelimb: A657T: 5.2 ± 0.31 cm, n = 20; WT: 5.7 ± 0.44 cm, n = 27, p<0.0001; T660K: 5.4 ± 0.56 cm, n = 20; WT: 5.9 ± 0.43 cm, n = 19, p=0.0032; hindlimb: A657T: 5.2 ± 0.33 cm, n = 20; WT: 5.7 ± 0.45 cm, n = 27, p<0.0001; T660K: 5.4 ± 0.50 cm, n = 20; WT: 5.9 ± 0.40 cm, n = 19, p=0.0053; Fig. 6E-G), as well as a reciprocal increase in stride frequency (not shown). In addition to this impairment in efficiency, *Grik2* mutant animals demonstrated a reduction in hindpaw angle (A657T: 15 ± 3.3°, n = 20; WT: 18 ± 3.2°, n= 27, p=0.0054; T660K: 12 ± 3.1°, n = 20; WT: 16 ± 2.9°, n = 19, p=0.0008; Fig 6H, I). This phenotype is noteworthy given that the opposite phenotype – an increase in hindpaw angle or splay - is commonly observed in mouse models of ataxia and neurodegeneration (de Haas et al., 2016; Guyenet et al., 2010; Sleat et al., 2004; Yerger et al., 2022). In T660K mice, a reduction in hindlimb stance width was observed in addition to reduced hindpaw angle compared to their WT littermates (T660K: 1.7 ± 0.32 cm, n = 21; WT: 1.9 ± 0.26 cm, n = 19, p=0.016) but was not observed in A657T mice (A657T: 2.1 ± 0.20 cm, n = 20; WT: 2.1 ± 0.15 cm, n = 27, p=0.34 ; Fig. 6H, J). We performed linear regression analysis and found only weak correlations between stride length and weight (A657T: R^2^=0.10, WT: R^2^=0.22, T660K: R^2^= 0.005, WT: R^2^= 0.14), suggesting that reduced mass was not responsible for smaller stride length in the mutant mice (data not shown).

### 3.7. Grik2 mutant mice display abnormal postures and dystonic-like movements

Gait alterations were explored further in a set of tests designed to characterize ataxic-like phenotypes caused by the *Grik2* mutation. We used a scale to assess ataxia in mice in which scores range from 0 to 3, with 0 indicating no deficits and 3 indicating severe ataxia (Guyenet et al., 2010). The overall score comprises the average results of three behavioral tests: kyphosis (abnormal spine curvature) (Fig. 7A), movement along a ledge (Fig. 7B), and hindlimb clasping (Fig. 7C). *Grik2* mutant mice displayed hunched and clasped postures and had more difficulty traversing a narrow ledge and consequently scored significantly higher (worse) than their WT littermates (kyphosis: A657T: 1.5 ± 0.5, n = 24; WT: 0.5 ± 0.4, n = 34, p<0.0001; T660K: 0.86 ± 0.68, n = 22; WT: 0.017 ± 0.091, n = 30, p<0.0001; ledge test: A657T: 1.0 ± 0.7, n = 24; WT: 0.3 ± 0.6, n = 34, p<0.0001; T660K: 1.9 ± 0.6, n = 22; WT: 1.1 ± 0.6, n = 30, p<0.0001; clasping: A657T: 2.0 ± 0.6, n = 24; WT: 0.2 ± 0.3, n = 34, p<0.0001; T660K: 2.3 ± 0.6, n = 22; WT: 0.0 ± 0.0, n = 30, p<0.0001; Fig. 7A-C). The average ataxia score for A657T and T660K mice were 1.5 ± 0.4 (n = 24) and 1.7 ± 0.4 (n = 22), respectively, consistent with a mild to moderate ataxic-like phenotype, whereas their WT littermates scored 0.3 ± 0.2 (n = 34, p<0.0001) and 0.4 ± 0.2 (n = 30, p<0.0001; Fig. 7D). This result and the gait impairments observed in the previous set of experiments show that the mutant KAR mice have a phenotype qualitatively similar to those displayed by other mouse models of ataxia (Cendelin, 2014; Guyenet et al., 2010).

**Figure 7.**
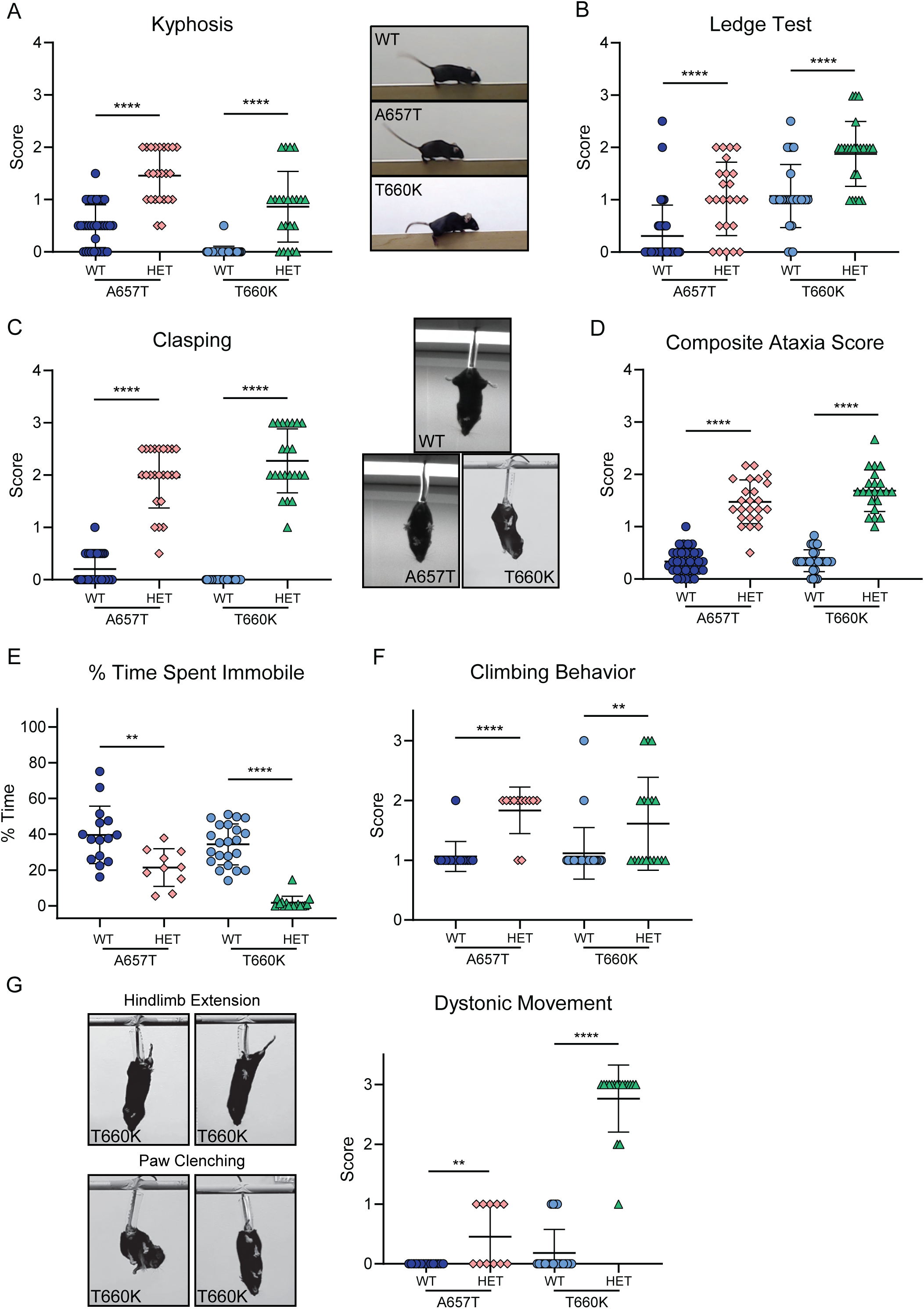
Grik2 GoF mice present with postures and movements associated with ataxic and dystonic-like phenotypes. A) Kyphosis scores demonstrated that both A657T and T660K mice had a greater degree of spinal curvature than WTs as shown in the representative images. Scale: 0: no kyphosis, 1: mild kyphosis, able to straighten spine, 2: persistent mild kyphosis, 3: persistent severe kyphosis. B) *Grik2* GoF mice had compromised movement when walking along a narrow ledge. . Scale: 0: ledge walking, dismount, 1: paw slips, 2: reduced hindpaw use, scooting, near fall, 3: immobility, fall. C) Paw clasping was scored during tail suspension. Both A657T and T660K mice showed significantly more clasping than WTs as shown in the representative images. 0: consistently splayed, 1: one limb retracted, 2: both limbs retracted, 3: both limbs entirely retracted, touching abdomen. D) A composite ataxia score was calculated by averaging the scores from the kyphosis, ledge, and clasping indices. *Grik2* GoF mice scored higher than their WT littermates. E) The percent time spent immobile during tail suspension was measured to quantify the rapid kicking and writhing-like behavior observed in *Grik2* GoF mice during tail suspension. Both A657T and T660K mice spent less time immobile. F) Escape behavior was assessed by measuring climbing attempts during tail suspension. *Grik2* GoF mice scored higher in this index, indicating that they were less likely to reach their tails. Scale: 1: able to curl/climb up tail to right self, 2: attempts to curl up and right self unsuccessfully, 3: no attempt to curl up. G) *Grik2* GoF mice scored higher for dystonic-like movements calculated based on clasping, aberrant hindlimb extension, and paw clenching analyses. Images show representative hindlimb extension and paw clenching in T660K mice. Scale: 0: no dystonic-like behaviors, 1: dystonic-like <50% of time, 2: dystonic-like >50% of time, 3: dystonic-like 100% of time. Asterisks indicate p-values: *, p < 0.05; **, p < 0.01; ***, p < 0.001; ****, p < 0.0001.

*Grik2* mutant mice were assessed for abnormal behaviors in addition to those associated with ataxia during the tail suspension test. During routine handling, both types of *Grik2* mutant mice were identifiable by their rapid and continuous kicking and writhing. We quantified this behavior and found that they spent markedly less time immobile during suspension compared to WT mice (percent time immobile: A657T, 21 ± 11 %, n = 10; WT, 40 ± 16 %, n = 15, p=0.0045; T660K, 1.7 ± 3.6 %, n = 17; WT, 34 ± 11 %, n = 22, p<0.0001; Fig. 7E). This distinctive kinetic movement also occurs in mice with deficits in proprioception due to conditional deletion of the Piezo2 mechanoreceptor (Woo et al., 2015), and indeed several individuals with the

*GRIK2* p.Ala657Thr variant exhibit proprioceptive dysfunction. *Grik2* mutant mice also were less likely to right themselves to grab their tails during suspension, which has been characterized as a naturalistic escape behavior (A657T: 1.8 ± 0.39, n = 12; WT: 1.1 ± 0.25, n = 16, p<0.0001; T660K: 1.6 ± 0.78, n = 18; WT: 1.1 ± 0.43, n = 26, p=0.0076; Fig. 7F). Most animals that were unable to reach their tails actively attempted to right themselves but failed.

Finally, we assessed if *Grik2* mutant mice exhibited postures associated with dystonia, including hindlimb clasping, paw clenching, and sustained hindlimb hyperextension. The T660K mice demonstrated these phenotypes consistently while suspended by their tail, while A657T mice exhibited clasping sporadically (T660K: 2.8 ± 0.56, n = 17; WT: 0.18 ± 0.39, n = 22, p<0.0001; A657T: 0.45 ± 0.52, n = 11; WT: 0.0 ± 0.0, n = 15, p=0.0070; Fig. 7G). Overall, *Grik2* mutants demonstrate abnormal movements and postures during tail suspension that are consistent with underlying impairments related to disorders like dystonia or ataxia.

### 3.8. Grik2 GoF mice are impaired in a test of spatial working memory

Individuals with the *GRIK2* p.Ala657Thr variant have ID, whereas those with the p.Thr660Lys are so severely impaired that they are unable to be assessed for cognitive function. We therefore tested *Grik2* mutant mice for deficits in short- and long-term memory tasks. Spatial working memory was tested in a variable DNMS alternating T-maze task. The training period requires that the mice first reach ≥70% correct alternating choices for at least three consecutive days of trials after habituation and pre-training consisting of forced alternations (Fig. 8A). In preparation for this task, mice are typically food-restricted to 85% of their normal body weight to enhance motivation to obtain the reward for correct choices. T660K mice, however, were acutely sensitive to food restriction and 83% of T660K mice in the cohort died during this initial pre-training period. We therefore limited food consumption for these mice and their littermates to only two hours before training and testing. During the training phase, A657T mice reached criteria in the same number of days as their WT littermates (A657T: 4.3 ± 1.5 days, n = 12; WT: 3.4 ± 0.7, n = 12, p=0.09), but T660K mice took almost 2 days longer on average (T660K: 5.4 ± 2.0, n = 9; WT: 3.5 ± 1.0, n = 11, p=0.02; Fig. 8B). Additionally, 31% (4 of 13) of T660K mice were not able to reach criteria, while all A657T mice were successfully trained.

**Figure 8.**
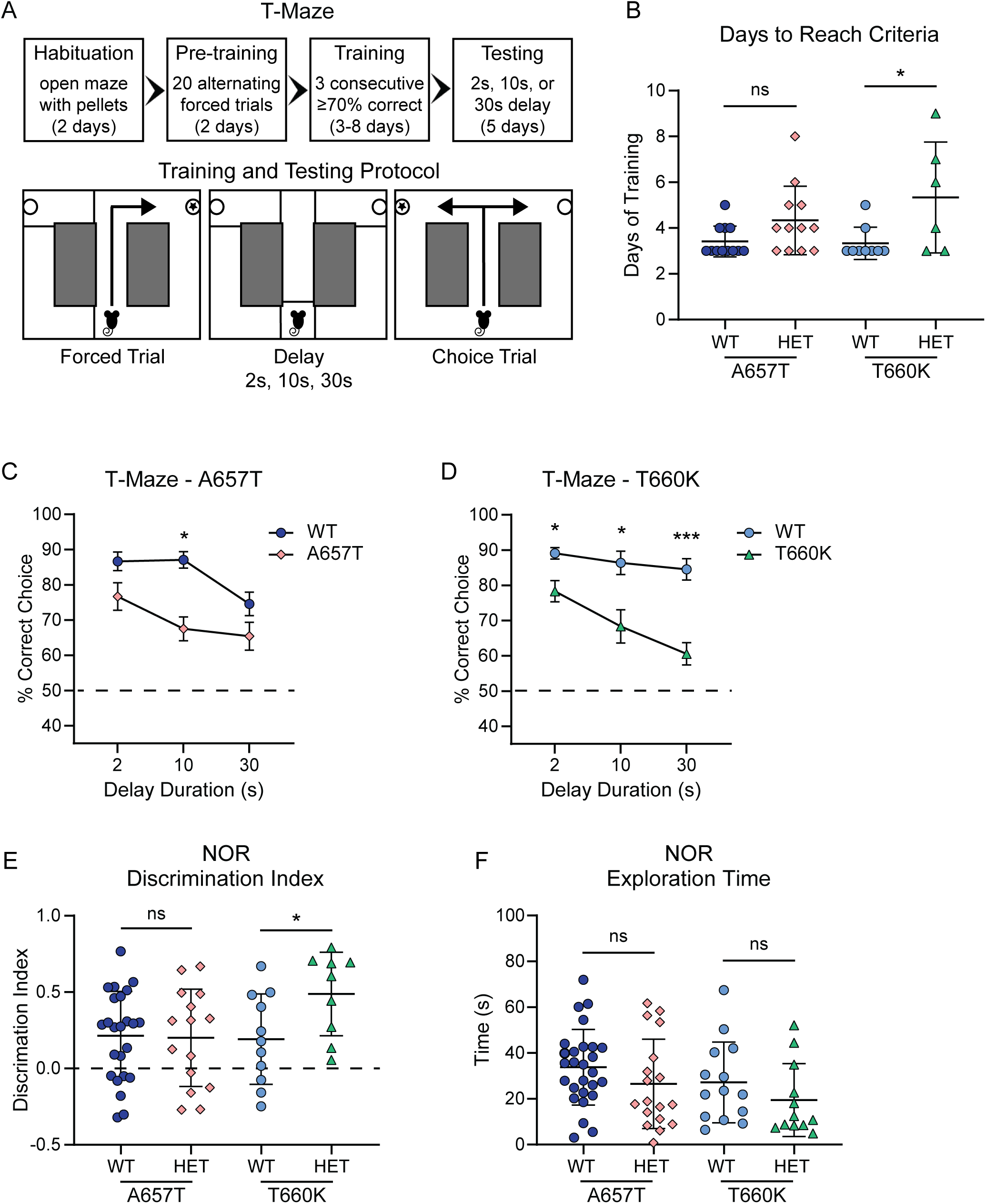
*Grik2* GoF mice have deficits in a test of working spatial memory but not in long-term recognition memory. A) Schematic of T-Maze testing. Top: Training proceeded as follows; 2 days of habituation, 2 days of forced-alternation pre-training, and then DNMS training until animals reach criterion of 70% correct choices. Testing then proceeds with variable delays over 5 days. Bottom: Schematic of each DNMS trial consisting of a forced trial, a period of delay, and a choice trial. B) The number of days to reach the criterion was equivalent for WT and A657T mice. In contrast, T660K mice took longer on average to reach the criteria. C) A657T mice performed worse than WT littermates when a 10 sec delay was imposed in the DNMS test. There was no difference in performance at a longer delay of 30 sec. D) T660K mice made more incorrect choices than WT mice at all three delays of 2, 10, and 30 sec in the DNMS testing. E) A657T animals showed the same level of novel object discrimination as their WT littermates. T660K mice displayed increased discrimination, indicating a higher level of interest in the novel object. F) Exploration time during the NOR test did not reveal any differences between the genotypes. Asterisks indicate p-values: *, p < 0.05; **, p < 0.01; ***, p < 0.001; ****, p < 0.0001.

After the mice reached the criterion, we tested them for alternation in a choice trial with a 2, 10, or 30 second delay. For GluK2(A657T) mice, we found a significant effect of genotype (F_1,22_ = 20, p=0.0002) that accounted for a larger percentage of performance variance than the delay itself (F_1.87,41.2_ = 6.8, p=0.0034). When comparing performance at each delay interval, A657T mice performed significantly worse than their WT littermates at a delay of 10 seconds (A657T: 68 ± 12%, n = 12; WT: 87 ± 8%, n = 12, p=0.0004; Fig. 8C). For T660K mice, genotype (F_1,18_ = 34.7, p<0.0001) again accounted for a larger percent of performance variance than delay (F_1.54,27.8_ = 7.2, p=0.0055). Furthermore, T660K mice performed significantly worse than their WT littermates at all 3 delays (2 s: T660K: 78 ± 9%, n = 9; WT: 89 ± 5%, n = 11, p=0.024; 10 s: T660K: 68 ± 14%, n = 9; WT: 86 ± 11%, n = 11, p=0.021; 30 s: T660K: 61 ± 10%, n = 9; WT: 85 ± 10%, n = 11, p=0.001; Fig. 8D). Thus, *Grik2* GoF mice have impaired spatial working memory.

We tested long-term memory in a NOR task, which assesses the relative proportion of time that the mice spend investigating a new object introduced into the testing cage. A657T mice did not display differences in novel object discrimination or in the time spent exploring both objects compared to their WT littermates (discrimination index: A657T: 0.20 ± 0.32, n = 15; WT: 0.21 ± 0.29, n = 24, p=0.89; exploration time: A657T: 27 ± 20 s, n = 18; WT: 34 ± 17 s, n = 27, p=0.11; Fig. 8E). T660K mice instead displayed increased novel object discrimination while exploring the objects the same amount as their WT littermates (discrimination index: T660K: 0.48 ± 0.27, n = 9; WT: 0.19 ± 0.30, n = 11, p=0.034; exploration time: T660K: 19 ± 16 s, n = 12; WT: 27 ± 18 s, n = 14, p=0.19; Fig. 8F). Thus, we conclude from these tests that the A657T and T660K mice exhibit a deficit in working but not long-term memory.

### 3.9. Neonatal Grik2 GoF mice have motor and vocalization impairments

*Grik2* expression in rodents is present across numerous brain regions as early as the second week of embryonic development and persists throughout adulthood (Bahn et al., 1994). The mutation in *Grik2* therefore is likely to alter KAR signaling and proper CNS development well before synapses are formed postnatally. We tested if the A657T and T660K mutations cause dysfunction in early stages of pup development in the form of aberrant motor or isolation-induced vocalizations.

Parameters of motor function in *Grik2* mutant and WT pups were assessed at three key developmental time points – P3, P7, and P10. We measured righting reflex, ambulation, hindlimb suspension, and forelimb suspension in tests adapted from a battery developed for a cerebral palsy mouse model (Feather-Schussler and Ferguson, 2016). Gait development is normal in A657T mice (genotype effect: F_1, 23_ = 0.0621, P=0.81; Fig. 9A), but T660K animals show a delay in walking development with many not demonstrating spontaneous walking during testing until P10 (genotype effect: F_1, 24_ = 39.2, p<0.0001; gait scores: P3: T660K: 0.46 ± 0.52, n = 13; WT: 0.92 ± 0.28, n = 13, p=0.033; P7: T660K: 0 ± 0, n = 13; WT: 1.8 ± 0.73, n = 13, p<0.0001; P10: T660K: 1.6 ± 1.3, n = 13; WT: 2.4 ± 0.65, n = 13, p=0.19; Fig. 9A). A657T pups again exhibit normal righting reflex development (genotype effect: F_1, 23_ = 2.84, p=0.11; Fig. 9B), while T660K mice exhibit a delay in developing this reflex at P3 that normalizes by P7 (genotype effect: F_1, 37_ = 25.9, p<0.0001; P3: T660K: 55 ± 8.9 s, n = 13; WT: 33 ± 19 s, n = 26, p<0.0001; P7: T660K: 15 ± 18 s, n = 13; WT: 5.2 ± 5.3 s, n = 26, p=0.19; P10: T660K: 2.0 ± 1.7 s, n = 13; WT: 1.1 ± 0.25 s, n = 26, p=0.28; Fig. 9B).

**Figure 9.**
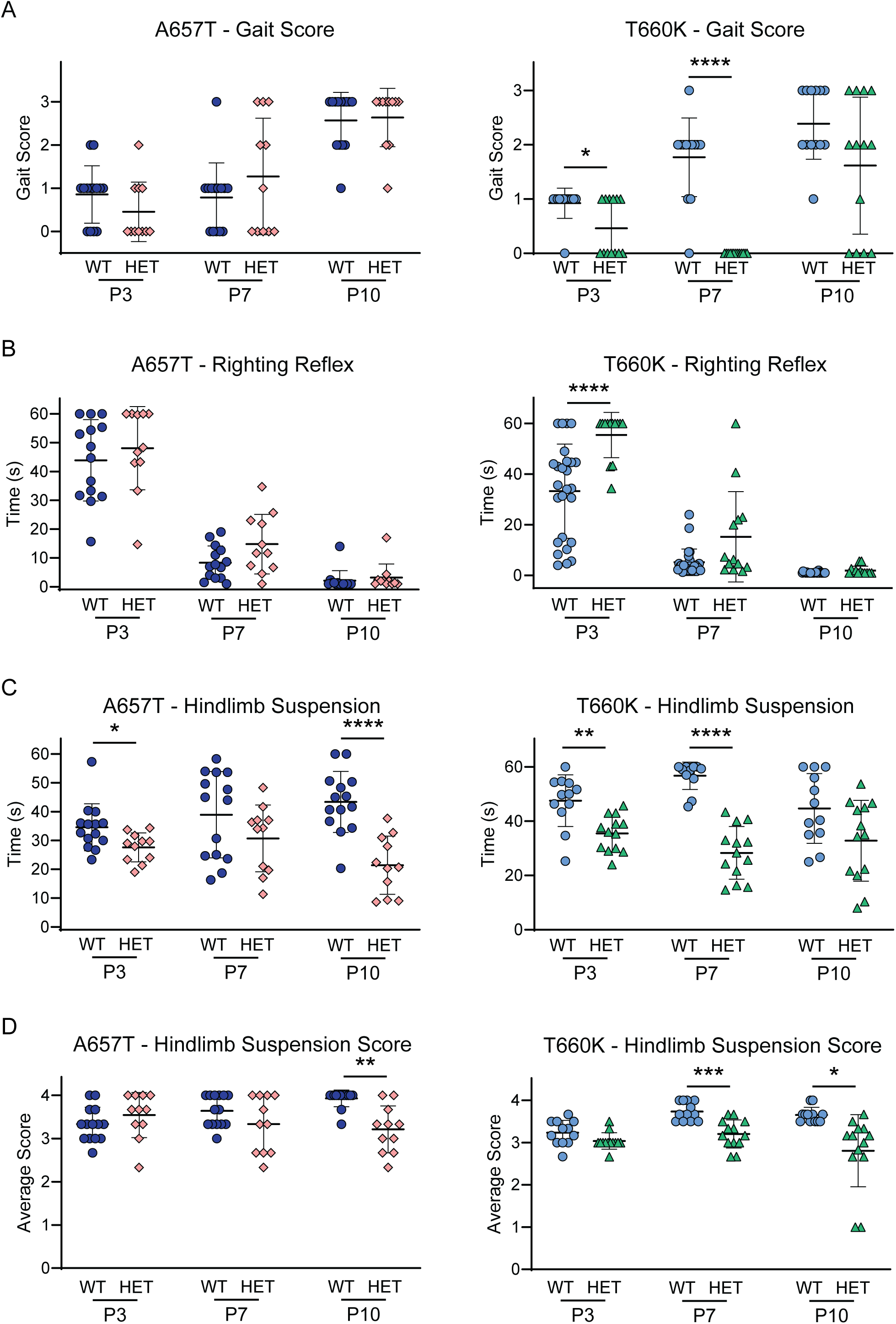
Deficits in motor behavior in *Grik2* GoF pups. A) There was no difference in gait scores for A657T mice compared to WT littermates. T660K mice scored lower at P3 and P7, indicating slower motor development. B) A657T mice showed normal development of a righting reflex, whereas T660K mice took longer than their WT littermates at P3. Scale: 0: no movement, 1: crawling with asymmetric limb movement, 2: slow crawling with symmetric limb movement, 3: fast crawling/walking. C) The length of time that pups remained suspended by their hindlimbs was measured to assess muscle strength. A657T pups stayed suspended for less time than WTs at both P3 and P10. T660K pups suspended for less time at both P3 and P7. D) A657T hindlimb posture was scored lower than WTs at P10. T660Ks scored lower at both P7 and P10. Scale: 4: normal separation, 3: less separation with infrequent touching, 2: minimal separation, frequent touching between hindlimbs, 1: clasping the majority of the time, and 0: constant clasping and/or fall. Asterisks indicate p-values: *, p < 0.05; **, p < 0.01; ***, p < 0.001; ****, p < 0.0001.

We tested limb strength and posture by suspending the mice from their hindlimbs, measuring the time to drop and relative angle of the hindpaws (Fig. 9C, D). The time to fall was significantly shorter for both A657T and T660K mice at the earliest postnatal age tested, P3, and more markedly different at P7 for T660K mice and P10 for A657T mice (A657T genotype effect: F_1, 23_ = 14.5, p=0.0009; P3: A657T: 28 ± 5.0 s, n = 11; WT: 35 ± 8.2 s, n = 14, p=0.046; P10: A657T: 21 ± 10 s, n = 11; WT: 43 ± 11 s, n = 14, p<0.0001; T660K genotype effect: F_1, 23_ = 50.6, p<0.0001 ; P3: T660K: 35 ± 6.6 s, n = 13; WT: 47 ± 9.5 s, n = 12, p=0.0049; P7: T660K: 28 ± 9.7 s, n = 13; WT: 57 ± 5.0 s, n = 12, p<0.0001; Fig. 9C), suggesting that muscle development is slowed in neonates. In addition, P10 A657T pups and both P7 and P10 T660K mice displayed altered posture during this test. Posture was scored on a scale of 0-4 based on proximal to distal placement hindlimb placement relative to each other and the animal’s abdomen (Feather-Schussler and Ferguson, 2016). A657T pups were not significantly different across the three ages tested (F_1, 23_ = 4.18, p=0.052), but hindlimb posture was scored significantly lower at P10 (A657T: 3.2 ± 0.54, n = 11; WT: 3.9 ± 0.19, n = 14, p=0.0039; Fig. 9D). T660K pups did display a significant effect of genotype (F_1, 23_ = 20.6, p=0.0001), with both P7 and P10 hindlimb posture scored significantly lower (P7: T660K: 3.2 ± 0.33, n = 13; WT: 3.7 ± 0.22, n = 12, p=0.0003; P10: T660K: 2.8 ± 0.85, n = 13; WT: 3.7 ± 0.18, n = 12, p=0.012; Fig. 9D). These results indicate that, as observed in adults (Fig. 7C), *Grik2* GoF mice hold their hindlimbs closer together, retracted toward their abdomens.

Neonatal mice transiently produce USVs as a means of communicating, particularly under stressful conditions like separation from their mothers (Ehret, 2005), with features like call rate and duration thought to influence maternal responsiveness (D’Amato et al., 2005; Ehret and Haack, 1981). USVs have been observed to be impaired in mouse models of NDDs with social deficits like ASD and are thought to model early communication difficulties experienced by people with delayed or impaired development of articulation or the use of language (Caruso et al., 2020), like those individuals with the *GRIK2* p.Ala657Thr variant. Children with the p.Thr660Lys variant are non-verbal (Stolz et al., 2021). We therefore tested *Grik2* mutant mice for developmental alterations in USVs in response to maternal separation. Mice were removed from their mothers individually and recorded for five minutes. The recordings were analyzed to quantify USV parameters including counts, duration, frequency bandwidth, and total syllable energy (Fig. 10A). Across these five parameters for A657T mice, a two-way ANOVA identified a significant effect of genotype in syllable duration (F_1, 33_ = 18.0, P=0.0002; Fig. 10C) and total syllable energy (F_1, 33_ = 16.3, P=0.0003; Fig. 10D). GluK2(A657T) mice generated fewer syllables at P3 (A657T: 48 ± 30 syllables, n = 16; WT: 140 ± 95 syllables, n = 19, p=0. 0016; Fig. 10B), suggesting that they are delayed in their development of ultrasonic call production. Syllable duration was significantly different at every time point (P3: A657T: 40 ± 11 ms, n = 16; WT: 51 ± 12 ms, n = 19, p=0.032; P7: A657T: 46 ± 9.1 ms, n = 16; WT: 56 ± 11 ms, n = 19, p=0.014; P10: A657T: 46 ± 8.7 ms, n = 16; WT: 56 ± 9.3 ms, n = 19, p=0.0097; Fig. 10C), indicating that A657T pups produce shorter USVs throughout these key developmental time points during maternal rearing. T660K mice similarly show deficits in USV production, but only in the number of syllables produced at both P3 and P7, normalizing by P10 (genotype effect: F_1, 28_ = 28.1, P<0.0001; syllable counts: P3: T660K: 51 ± 45, n = 13; WT: 160 ± 150, n = 17, p=0.023; P7: T660K: 94 ± 52, n = 13; WT: 330 ± 130, n = 17, p<0.0001; P10: T660K: 110 ± 62, n = 13; WT: 180 ± 120, n = 17, p=0.12; Fig. 10B). *Grik2* GoF mice therefore exhibit robust behavioral phenotypes very early in development in both motor domains and in USVs, with some of those abnormalities resolving in the first postnatal week and other phenotypes emerging later in maturation.

**Figure 10.**
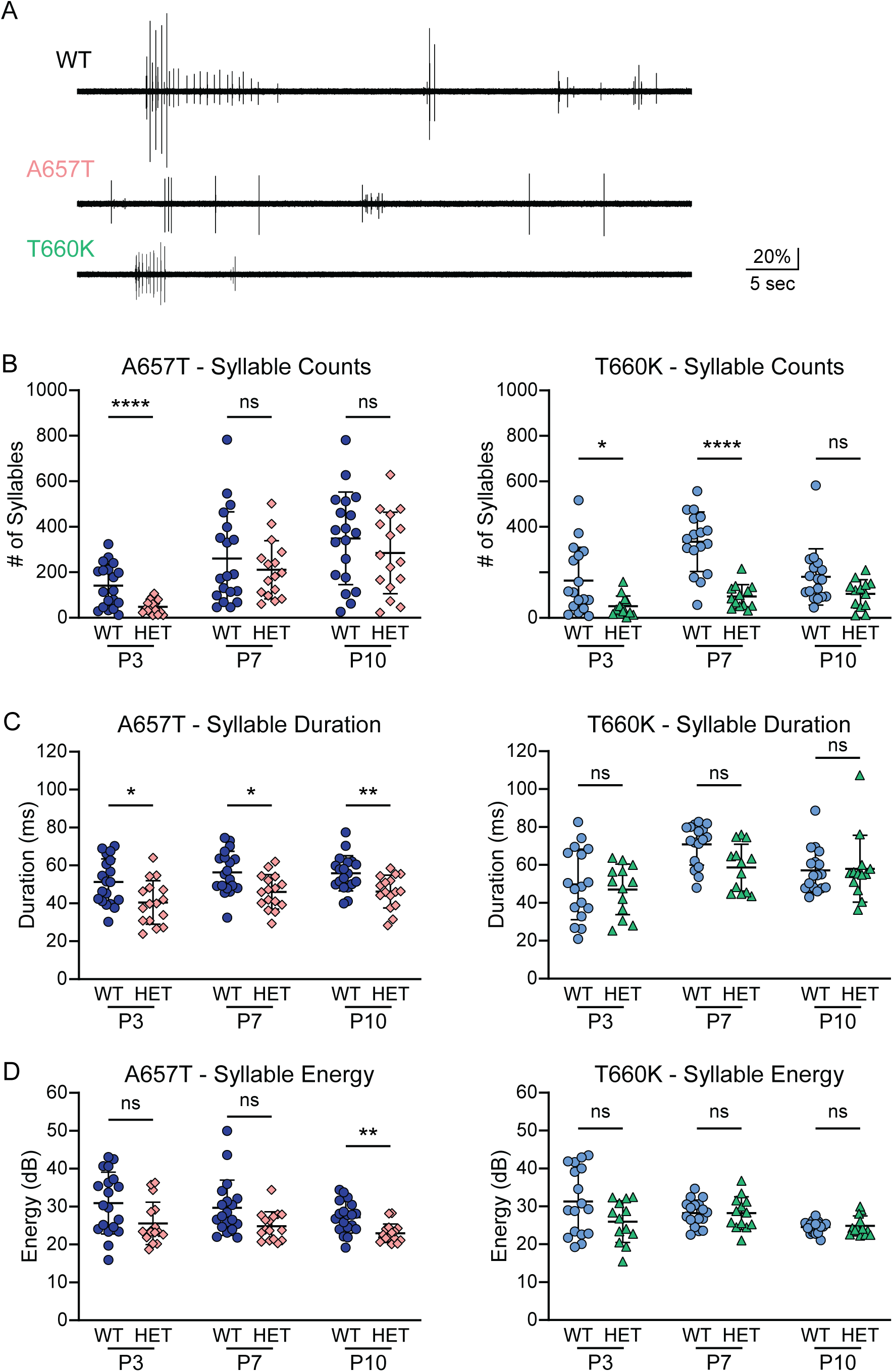
USVs during early development of *Grik2* GoF pups. A) Representative USV traces for WT, A657T, and T660K mice at P3. B) Syllable counts were measured to assess pup vocalization frequency. A657T mice produced fewer USVs at P3, and T660K mice produced fewer USVs at P3 and P7. C) A657T mice produced shorter vocalizations at all 3 timepoints. T660K mice had USVs of the same duration as WT. D) A657T mice produced weaker USVs at P10 as measured by syllable energy. T660K mice were again equivalent to WT. Asterisks indicate p-values: *, p < 0.05; **, p < 0.01; ****, p < 0.0001.

## 4. Discussion

Disruptions in *GRIK2* have long been associated with a range of neurodevelopmental and psychiatric disorders, including ID, epilepsy, and ASD (Hansen et al., 2021). However, the extent to which *de novo* variants underlie nonsyndromic NDDs was unknown until we identified the first pathogenic missense mutations in this KAR subunit gene in humans (Guzmán et al., 2017; Stolz et al., 2021). Individuals with those variants have disorders of varying degrees of severity with symptoms that include ID, GDD, impairments in the use of language, and, in a subset of patients, epilepsy (Guzmán et al., 2017; Stolz et al., 2021). In the current study, we characterized two *Grik2* knockin mouse models of the most common *GRIK2* variants, p.Ala657Thr and p.Thr660Lys, to better understand how *GRIK2* mutations can drive developmental dysfunction. The diverse behavioral phenotypes exhibited by the two mouse lines support classification of the *GRIK2* variants as autosomal dominant, monogenic causes of pathogenic NDDs. They also exhibit a comparative severity in phenotypes similar to those in children with the variants, underscoring the face validity of the mice as models of human disease. Moreover, our findings demonstrate that *Grik2* GoF variants produce symptom profiles that differ from and are more severe than those in mice with a LoF in *Grik2*. This work represents a critical step toward understanding how *GRIK2* GoF variants contribute to NDDs and lays the groundwork for future studies into the underlying mechanisms of KAR dysfunction in human disease.

### 4.1. Grik2 mutant mice partially recapitulate human GRIK2 p.Ala657Thr and p.Thr660Lys disorders

Our primary objectives of the current research included testing if mouse models supported the prediction that the *GRIK2* variants are causative for human NDDs and determining to what extent the mice reproduce core features of the diseases. Our results provide strong support for the damaging effects of these two *GRIK2* mutations and demonstrate that *Grik2* GoF mouse models exhibit diverse phenotypes analogous to many aspects of the human disorders. Moreover, the relative severity of the mouse phenotypes mimics that in individuals. That is, T660K mice exhibit more severe phenotypes than A657T across nearly all functional domains, consistent with the more debilitating clinical presentation associated with the p.Thr660Lys variant.

Dysfunction in the KAR mutant mice is apparent at early developmental stages. Viability to weaning age is reduced in both lines, but T660K mice are compromised to a greater degree in that only 25% of pups had the mutation instead of the expected 50% from heterozygous x WT breedings. A657T mice show a modest deviation from the expected ratio, comprising 43% of total offspring. Also, *Grik2* GoF mice weighed less than their WT littermates. T660K mice on average had body weights lower than their WT counterparts starting as early as postnatal day 10, whereas A657T mice do not show relative weight reductions until approximately five weeks of age. Both lines of mice exhibited abnormal pup behaviors during the first postnatal week, including impairments in motor coordination and USV production, but only T660K animals were delayed in maturation of righting reflexes and normal gait. These observations underscore the pathogenic nature of the *Grik2* mutations even before synaptogenesis and development of cortical circuits.

*Grik2* mutant mice differentially exhibit other phenotypes analogous to those in humans. Susceptibility to seizures is a striking example. All three individuals with the *GRIK2* p.Thr660Lys variant have seizure disorders, and, similarly, by late adolescence T660K mice begin to display handling-induced seizures that increase in frequency into early adulthood. In contrast, the p.Ala657Thr variant does not cause epilepsy and seizures in the GluK2(A657T) mice are infrequent. The *Grik2* mutant mice also exhibit cognitive impairments with phenotypic differences that align with the severity of the analogous human variants. The p.Ala657Thr variant is associated with moderate to severe ID, while p.Thr660Lys results in profound cognitive impairment and minimal responsiveness to external stimuli. *Grik2* GoF mice both exhibit spatial working memory deficits in the DNMS task; moreover, 31% of T660K mice failed to reach the criterion of 70% correct choices. In contrast, A657T and T660K mice discriminate novel objects to the same degree as their WT counterparts, suggesting that they retain long-term recognition memory.

Some aspects of motor deficits observed in humans are mirrored in the mouse phenotypes. Individuals with the p.Ala657Thr variant are ambulatory but exhibit varying degrees of movement and proprioceptive dysfunction that manifest in part as ataxia, imbalance, and uncoordinated gait in the nine affected individuals. In contrast, children with the p.Thr660Lys variant cannot walk, have a variety of muscle tone abnormalities and are limited in their ability to support themselves without assistance. The analogous mouse lines show gait and balance impairments, kyphosis, and hindlimb clasping, which are established markers of ataxia and other motor disorders in rodents (Brooks and Dunnett, 2009; Guyenet et al., 2010).

Recently, one individual with the p.Ala657Thr variant was diagnosed with dystonia based on increasingly contorted postures of her hands and feet (Swanson, personal communication). Dystonic symptoms appear in some but not all monogenic mouse models as abnormal clasping or postures and twisting motions when suspended (Liang et al., 2014; Pappas et al., 2015; Shashidharan et al., 2005), as we observed in both A657T and T660K mice, but these phenotypes are associated with other NDDs as well (Oleas et al., 2013). Definitive classification of these movements as dystonic in the KAR mutant mice will require further analysis using electromyography (EMG) or related assessments (Jinnah et al., 2005; Pocratsky et al., 2023).

### 4.2. GoF in KAR signaling causes distinct and more severe phenotypes than LoF Grik2

GoF variants in mice produce phenotypes that are distinct from and, in some cases, the antithesis of those observed in *Grik2* knockout (KO) models. Homozygous *Grik2* KOs are viable and display increased activity and naturalistic behaviors (Shaltiel et al., 2008; Xu et al., 2017), (but see Iida et al., 2021), improvements in spatial learning (Micheau et al., 2014; Mulle et al., 1998), mild impairments in balance, and increased time spent immobile during tail suspension (Iida et al., 2021). No abnormal phenotypes have been reported in heterozygous *Grik2* KOs. In contrast, heterozygous *Grik2* GoF mice show attenuated innate behaviors, decreased spatial memory task performance, impaired motor performance across multiple domains, and decreased time spent immobile during tail suspension. Not surprisingly, KO mice are resistant to KA-induced seizures (Mulle et al., 1998), whereas both A657T and T660K mice exhibit heightened seizure susceptibility and lethality following exposure. This dichotomy in seizure phenotype represents a particularly robust functional divergence and underscores the role of GluK2-containing KARs in induction of seizures with this potent chemoconvulsant (Baudouin et al., 2024; Boileau et al., 2023; Crepel and Mulle, 2015).

Comparative findings from both mouse models and human genetic data further support the conclusion that GoF variants in GluK2 genes are more deleterious to neural development than LoF. Unlike *Grik2* KO mice, homozygous A657T and T660K mice are not viable. This mirrors human genetic observations: while GoF variants are *de novo* and heterozygous, suspected LoF variants can be inherited, and some individuals carry homozygous LoF alleles (Córdoba et al., 2015; Motazacker et al., 2007). A heterozygous, *de novo GRIK2* variant predicted to cause a LoF was reported in one individual diagnosed with ASD but no other symptoms, aligning with the lack of phenotypic observations in *Grik2* heterozygous mice (Stolz et al., 2021). Additionally, humans with homozygous *GRIK2* deletions may exhibit mild to severe ID, but they do not all have seizures (Córdoba et al., 2015; Motazacker et al., 2007) - in contrast to the often intractable, mixed seizure disorders observed in individuals with the p.Thr660Lys GoF variant (Stolz et al., 2021). Motor deficits are also less pronounced or absent in both KO mice (Iida et al., 2021) and humans with LoF mutations (Córdoba et al., 2015; Motazacker et al., 2007; Stolz et al., 2021), whereas both humans and mice with GoF variants have prominent motor deficits. Similar asymmetric impacts of GoF versus LoF mutations have been reported in other ion channelopathies, like in *GRIA3* and *SCN2A*, where GoF variants lead to early-onset epilepsy and movement disorders, while LoF is more often associated with milder or later-onset (Rinaldi et al., 2024; Sanders et al., 2018). These findings therefore reinforce the importance of variant-specific knock-in models for understanding *GRIK2*-linked pathophysiology.

One potential explanation for the divergence in phenotypes between LoF and GoF models is that compensatory mechanisms - such as upregulation of functionally similar subunits or changes in synaptic plasticity - can partially mitigate the consequences of LoF, whereas GoF effects are less amenable to homeostatic correction. This imbalance might result from ion permeability properties of KARs containing GluK2 subunits; *Grik2* mRNAs harbor a Q/R site at which ADAR2- mediated RNA editing converts a genomically encoded glutamine (Q) to arginine (R), thereby reducing the calcium permeability of KARs containing these subunits. During early development, *Grik2* transcripts are predominantly unedited and therefore encode calcium-permeable channels, but by adulthood approximately 90% of *Grik2* transcripts are edited, making GluK2-containing receptors calcium-impermeable (Bernard et al., 1999; Paschen et al., 1997). Excessive calcium influx driven by GoF variants may be especially disruptive during early development, when calcium signaling is critical for neurodevelopmental processes such as activity-dependent gene expression, migration, and synaptogenesis (reviewed in Rosenberg and Spitzer, 2011). This idea is consistent with prior observations that loss of ADAR2-mediated RNA editing (resulting in exclusively calcium-permeable KARs) is lethal in mice due to seizure susceptibility (Higuchi et al., 2000).

### 4.3. Potential underlying neural causes of Grik2 GoF phenotypes

The presence of behavioral deficits in *Grik2* GoF mice and symptoms in humans with *GRIK2* GoF variants across domains suggests that neurological deficits may be widespread across brain regions. *Grik2* expression begins about a week before birth in rodents and is widespread across brain regions throughout development and into adulthood (Bahn et al., 1994; Bischoff et al., 1997; Cui et al., 2012; Wisden and Seeburg, 1993; Zhang et al., 1996), with similar widespread *GRIK2* expression observed in adult humans (Porter et al., 1997). It is possible that structural abnormalities contribute to pathology in children with the p.Thr660Lys variant, because their MRIs revealed white matter abnormalities and reductions in neuropil volume (Guzmán et al., 2017; Stolz et al., 2021). In contrast, MRIs from individuals with the *GRIK2* p.Ala657Thr variant show intact myelination and no signs of neurodegeneration (Guzmán et al., 2017; Stolz et al., 2021). Gross brain morphology in *Grik2* GoF mice is not obviously altered compared to their WT littermates in the cortical, striatal hippocampal, or cerebellar regions pictured in Figure 1C, but finer aberrations in morphology might be revealed with higher resolution analyses.

*Grik2* GoF mice display marked reductions in naturalistic behaviors - including exploration, rearing, grooming, digging, and nestlet shredding. The striatum plays a central role in initiating and coordinating internally motivated behaviors like digging and grooming (Lv et al., 2025; Marshall et al., 2024; Minkowicz et al., 2023; Puighermanal et al., 2020), and grooming, in particular, is a well-characterized, stereotyped behavior modulated by the striatum, neocortex, and brainstem (Kalueff et al., 2016). While naturalistic behaviors are often increased or repetitive in mouse models of NDDs, the reduced performance seen in *Grik2* GoF mice more closely resemble phenotypes observed in models of basal ganglia disorders such as Parkinson’s and Huntington’s disease (reviewed in Kalueff et al., 2016). These findings indicate that the widespread naturalistic behavioral reductions observed in *Grik2* GoF mice might arise from disruptions in striatal circuits, consistent with *Grik2* expression and the function of GluK2-containing KARs in mediating synaptic transmission and short-term plasticity at cortico-striatal synapses (Marshall et al., 2018; Xu et al., 2017) and modulating GABAergic signaling onto spiny projection neurons (Chergui et al., 2000).

*Grik2* GoF mice and humans with the analogous variants exhibit motor impairments that suggest dysfunction across multiple motor circuits. Ataxia is most commonly associated with cerebellar pathology (reviewed in Cendelin, 2014), but similar motor deficits and abnormal postures, such as hindlimb clasping and kyphosis, have also been reported in models of basal ganglia or neocortical dysfunction (Fernagut et al., 2002; Lalonde and Strazielle, 2011). Reduced stride length is observed in models of ataxia but is also a hallmark of Parkinsonian and Huntington gaits (Blin et al., 1990; Koller and Trimble, 1985). Other mouse models that exhibit this phenotype include those of striatal dysfunction (Amende et al., 2005), neurodegeneration (da Conceicao Pereira et al., 2021; Preisig et al., 2016), and various NDDs (Gadalla et al., 2014; Matas et al., 2021). Beginning in early development, *Grik2* GoF mice display altered posture that persists into adulthood in the form of both reduced hindlimb distance during tail suspension and reduced hindlimb angle during gait.

This phenotype contrasts with the increased hindlimb angle - or “duck feet” posture - commonly seen in ataxic models (Guyenet et al., 2010), but resembles the decreased angle observed in ALS models (Mancuso et al., 2011). Together, these phenotypes implicate cerebellar, basal ganglia, cortical, and spinal cord dysfunction - all prospective sites of disrupted GluK2-containing KAR function - and suggest that *Grik2* GoF-related motor impairments may arise from a complex interplay across multiple motor pathways.

Increased propensity for seizures in *Grik2* GoF mice and spontaneous seizures in T660Ks can originate from numerous brain regions (Chauhan et al., 2022), and their underlying pathophysiology can include ion channel dysfunction, altered excitatory/inhibitory balance, and aberrant network synchronization (Gonzalez et al., 2019). Upregulation of GluK2-containing KARs – specifically, aberrant synaptic incorporation – contributes to seizure generation in TLE models (reviewed in Mulle and Crepel, 2021). Previous work demonstrated that GluK2(A657T) subunits are incorporated into functional synaptic KARs at hippocampal mossy fiber synapses, which exhibit a prolonged EPSC_KA_ decay and constitutive currents consistent with their GoF activity (Nomura et al., 2023). Consequently, CA3 pyramidal neurons in mutant mice exhibit a higher frequency of spontaneous spike firing that could contribute to enhanced seizure susceptibility. The KA-induced seizure hypersensitivity and abnormal cortical activity observed in T660K mice indicates that this mutant subunit is also expressed and incorporated into functional receptors and demonstrates that GluK2(T660K) subunits do not have a fully dominant-negative effect on KAR signaling. EEGs in T660K mice contained sub-ictal aberrant firing across cortical and hippocampal regions, but it remains to be determined how their spontaneous seizures may arise. Notably, seizure attenuation via miRNA-mediated suppression of GluK2-encoding genes has shown promise in preclinical models and is now being tested in clinical trials (Baudouin et al., 2024; Boileau et al., 2023). This miRNA-based therapy, along with other genetic therapies, may offer effective treatment for seizures and potentially additional core symptoms of *GRIK2* disorders.

### 4.4. Conclusion

In this study, we found that knock-in mouse models of the two most common pathogenic human *GRIK2* GoF variants – p.Ala657Thr and p.Thr660Lys - exhibit behavioral dysfunction across multiple domains, overlapping with symptoms in the analogous human disorders. Our results demonstrate that heterozygous GoF in GluK2-encoding genes can cause diverse, developmentally emergent deficits, consistent with autosomal dominant, monogenic etiology. The A657T and T660K mouse phenotypes mirror the relative severity seen in humans and support the conclusion that GoF variants produce more severe and distinctive outcomes from LoF, underscoring the value of variant-specific models for studying *GRIK2*-linked pathophysiology. As sequencing becomes more accessible, more individuals will be identified with pathogenic *GRIK2* variants. Our *Grik2* GoF mouse models provide powerful preclinical platforms for testing novel therapies that directly target genetic etiologies including knockdown and editing strategies. Furthermore, future investigation into the underlying neural correlates may elucidate pharmacological targets capable of mitigating the core symptoms of these rare but debilitating disorders, in addition to enhancing our understanding of the role of KARs in neurodevelopment and disease.

## Acknowledgements

This work was supported by National Institutes of Health/National Institute of Neurological Disorders and Stroke R01NS105502 and the National Institutes of Health/National Eye Institute R01EY032506 to G.T.S, as well as the National Institutes of Health/National Institute of Neurological Disorders and Stroke T32NS041234 to B.T.W. G.T.S. also thanks the Murphy family for additional support for these studies. We thank Drs. John Armstrong, Anis Contractor, Nicole Hawkins, Jennifer Kearney, John Marshall, Jones Parker, and Peter Penzes at Northwestern University for their technical and material assistance. The genetically engineered mice were generated with the assistance of Northwestern University Transgenic and Targeted Mutagenesis Laboratory. This work also was supported by the Northwestern University Behavioral Phenotyping Core.

## Notes

### Competing Interest Statement

The authors have declared no competing interest.

